# Electrically Conductive DNA-Inspired Coating for Intracortical Neural Microelectrodes

**DOI:** 10.1101/2023.06.19.545632

**Authors:** Ian Sands, Alpaslan Ersöz, Wuxia Zhang, Libo Zhou, Will Linthicum, Sabato Santaniello, Bryan Huey, Martin Han, Yupeng Chen

**Author notes:** Corresponding authors: Dr. Martin Han Dr. Yupeng Chen.

## Abstract

Different from conventional electrodes, intracortical neural microelectrodes are the size of one or several neural cells. Due to the limited space for cell-electrode connections, the bio-integration between each neural cell and the electrode surface is critical. To improve bio-integration and electrode functions, various coating materials (such as conductive polymers (CPs), carbon nanotubes (CNTs), and natural hydrogels) have been developed aiming to provide an enhanced interface for neuron recruitment, bio-anchorage, and electrical function. However, synthetic materials usually have limited biocompatibility and/or relatively high cytotoxicity, while biological materials present poor electrical functions. Therefore, current coatings possess biological, functional or electrochemical limitations that are not optimal for intracortical neural microelectrodes. To overcome this obstacle, we developed an electrically conductive coating based on biological molecules, named Janus base nano-coating (JBNc). JBNc is formed by Janus base nanotubes (JBNts) which are a family of nanotubes assembled from engineered DNA base pair units. Based on the long-distance translocation ability of the π electrons of JBNts, we developed them into an electrically conductive coating on the electrode surface. For the first time, we reported the DNA-inspired JBNc had an electrochemical performance that met and exceeded standard metal electrode surface in cyclic voltammetry, impedance spectroscopy, charge injection capacity tests, and neural recording. Moreover, we demonstrated enhanced bio-anchorage and microelectrode interface integration using SEM and AFM. Importantly, we demonstrated enhanced functional response to JBNc microelectrodes with immunohistochemical staining and RNA sequencing (RNAseq) analysis. Using cell viability assays, we also showed the benefits of DNA-mimicking chemistry of Janus base nanomaterials compared to conventional microelectrode coatings. We anticipate that these results will serve as a foundation for the continued development and study of JBNc to enhance interface dynamics and ultimately the performance and reliability of brain microelectrodes.

## Introduction

Since intracortical microelectrodes achieve high spatial resolution and single-neuron recording capabilities, their functional efficacy depends on successful integration with surrounding neural tissue. Their electrode geometry, similar to the size of single neurons, further stresses the importance of local bio-integration since the loss of neuron anchorage will greatly diminish accurate and consistent probe function. Successful integration between neural tissue and intracortical microelectrode is heavily dependent on the physical, biological and electrochemical properties governing the interface between them. A deficiency in one or more of these properties can significantly reduce the success of intracortical microelectrodes by discouraging healthy neuron attraction, which can contribute to poor long-term performance. To avoid chronic device isolation, it is essential to encourage bio-integration between neuron and foreign substrate, reinforcing the likelihood of efficient cell-to-electrode communication over an extended period. Conventional microelectrode materials (such as gold, iridium, platinum, etc.) present poor bio-integration with surrounding neurons, and may be susceptible to the unfavorable local environment conditions for neuron anchorage and function. This bio-incompatible or even cytotoxic microenvironment can induce apoptosis in otherwise healthy neurons, while simultaneously discouraging neuron integration and proper function on the microelectrode. A decreased healthy neuron attachment over time due to unfavorable substrates and cyclic tissue degradation is postulated to be partially responsible for many documented cases of diminishing robust electrical performance[1]. The encouragement of healthy cell bio-integration and anchorage to electrode sites is therefore essential for establishing a successful brain (intracortical) microelectrode interface that reduces the potential for long-term failure.

The poor bio-integration of conventional microelectrode materials is mainly due to their unfavorable physical and chemical properties such as stiffness and inorganic material composition, so a variety of coating materials have been developed to improve their surface properties. Polymer-based coatings such as polyethylene glycol (PEG) have been implemented and studied in an attempt to reduce probe stiffness and match the elastic modulus of the native tissue environment[2]. However, the chemical structure of some soft polymer coatings doesn’t allow for mediation of electron transport which may lead to suboptimal electrical performance. Reduced electron transfer caused by non-conductive polymer coatings has directed attention towards electrically conductive polymers (CPs), capable of increasing physical substrate favorability while maintaining the functional efficacy of the microelectrode. However, metal electrode materials like iridium oxide and gold still hold significant advantages over CP-based electrodes. CPs are less efficient for the transfer of electrons and are subsequently less electrically conductive compared to metal electrodes. With regards to microelectrode coatings, CPs such as Polypyrrole (PPy) have been implemented in many *in vivo* and *in vitro* applications, demonstrating their conductive capabilities [3]. CP coatings, however, present mixed biocompatibility profiles at higher concentrations with special consideration given to potential cytotoxicity caused by polymerization and doping techniques[4]. Many CPs claim sufficient biocompatibility regarding their short-term interactions with neural tissue. However, chronic stimulation of intracortical microelectrodes may release toxic byproducts resulting from chemical degradation. These byproducts are taken up by surrounding cells, introducing chronic cytotoxicity to the local tissue microenvironment. Conductive nanomaterial coatings like carbon nanotubes (CNTs) present excellent electrical conductivity potential, but present cytotoxic limitations to neuron/tissue cells similar to CPs, which limits their applications[5, 6]. Purification techniques, chemical structure, and natural oxidative stress imbalances can lead to significant local toxicity [7–10]. In an effort to reduce adverse biological response, natural hydrogels (such as chitosan, agarose, etc.) have gained attention in recent years due to their ability to meet electrical function criteria for microelectrode stimulation and recording[11, 12]. But natural hydrogels also possess practical disadvantages, including increased physical distance between electrode interface and target tissue. The conductivity of these hydrogels is also reliant on free-moving ions within their high water content. This high ion/water content eventually causes electrolysis which will change local pH and continue to generate toxic byproducts[13]. Fabrication techniques must also be taken into consideration, as the incorporation of CPs into non-conductive templates often lead to agglomeration of conductive species resulting in heterogeneous conductivity [10]. Existing coating methods present biocompatibility concerns, fabrication complications and electrical conductivity interference. In particular, while some coatings offer reduced impedances which reflect neural recording property, most have not demonstrated improvements in neural stimulation with relevant charge injection capacity [14–21]. Herein lies the incentive to develop a novel coating predicated on DNA-base nanomaterials that may overcome complications associated with current coating techniques.

JBNts are a family of Janus base nanomaterials (JBNs) self-assembled from G^C or A^T units mimicking DNA base pairs via hydrogen bonds and the base stacking effect[22]. The self-assembled DNA-mimicking structure of JBNts has demonstrated excellent bio-integration in terms of improved cell adhesion, proliferation and long-term functions with various types of tissue cells[23, 24]. Moreover, JBNts present significantly lower cytotoxicity than polymers and CNTs due to their non-covalent structures and biomimetic chemistry. All these findings have suggested promising potential as a biocompatible coating for neurological applications. JBNts utilize the assembly of DNA base units into an sp2 hybridized aromatic-ring system (rosette). Long distance delocalization of electrons is achieved following ν-ν base stacking of the rosettes comprised of those assembled base units. Interestingly, CNTs also present long-distance delocalization of electron through the π-π bond system. Because CNTs aromatic rings lie parallel to the length of the nanotube, their electrons can travel through the conjugated π bonds alongside the tube. Although similar to CNTs in shape, we discovered that JBNts aromatic rings lied on top of one another, allowing orthogonal p-orbitals to enable long distance electron dislocation between and through rosette stacks within the nanotube[25]. Based on the unique electrochemical property of JBNts, for the first time, we developed the JBNc which can effectively mobilize electrons while lying flat on the surface in a high density. Furthermore, JBNs are biomolecule-based materials self-assembled based on non-covalent hydrogen bonds, improving upon cytotoxic concerns with CNTs and CPs (such as PPy) which are synthetic materials formed by non-biodegradable C-C covalent bonds. Moreover, JBNc presents more robust electrical conductivity than natural hydrogels since their conductivity mechanism is not contingent on water and ion content[26]. Additionally, because the conductivity of JBNc relies on π electrons but not ions, it does not result in electrolysis (a common problem in hydrogels that can change pH and generate by-products)[13]. These key differences allow JBNc to create a new class of electrically conductive biomimetic coating. To our knowledge, JBNc is the first instance of an electrically conductive coating comprised of biological molecules.

In this study, we have developed controlled JBNt self-assembly and deposition resulting in the JBNc on the iridium and gold electrode surface as well as the silicon dioxide-based insulation material of intracortical microelectrodes. We conducted measurements to study JBNc capability of standalone charge conduction and impedance performance parameters on functional microelectrode probes. Fluorescence staining was performed to study cell recruitment to JBNc probes and proliferation assays were performed to study coating conditions. Additional immunohistochemical staining was used to analyze expression of β1 integrin, NeuN and NF-200 antigen production. Scanning Electron Microscopy (SEM) and Atomic Force Microscopy (AFM) was used to study integration and bio-anchorage at the interface between control and JBNc samples and differentiated neurons. Additionally, we conducted a concentration-based cytotoxicity assessment to demonstrate lack of cytotoxicity in comparison to other popular coating materials. Finally, we performed RNA sequencing (RNAseq) on SH-SY5Y neural cells grown on electrodes with and without JBNc to analyze their transcriptomic response following 2-week culture. This data serves as the benchmark for developing JBNc as an electrically conductive nano-coating for enhanced brain-to-electrode interfaces.

## Results

### Coating and Characterization

JBNt’s DNA-mimicking chemical structure is shown in **Figure 1**. Hydrogen bonding in **Figure1B** facilitates rosette assembly while closely stacked aromatic rings in **Figure 1C & D** enable π-π electron transfer. JBNt was prepared at a concentration of 1mg/mL and deposited onto the two shanks of intracortical microelectrodes. The microelectrode probe design used in this study is shown in **Figure 1** with the shanks constituting the major regions of interest. SEM and AFM imaging within the recording/stimulating electrode sites confirmed that the JBNt arrange horizontally on the microelectrodes (**Figure 1**).

**Figure 1.**
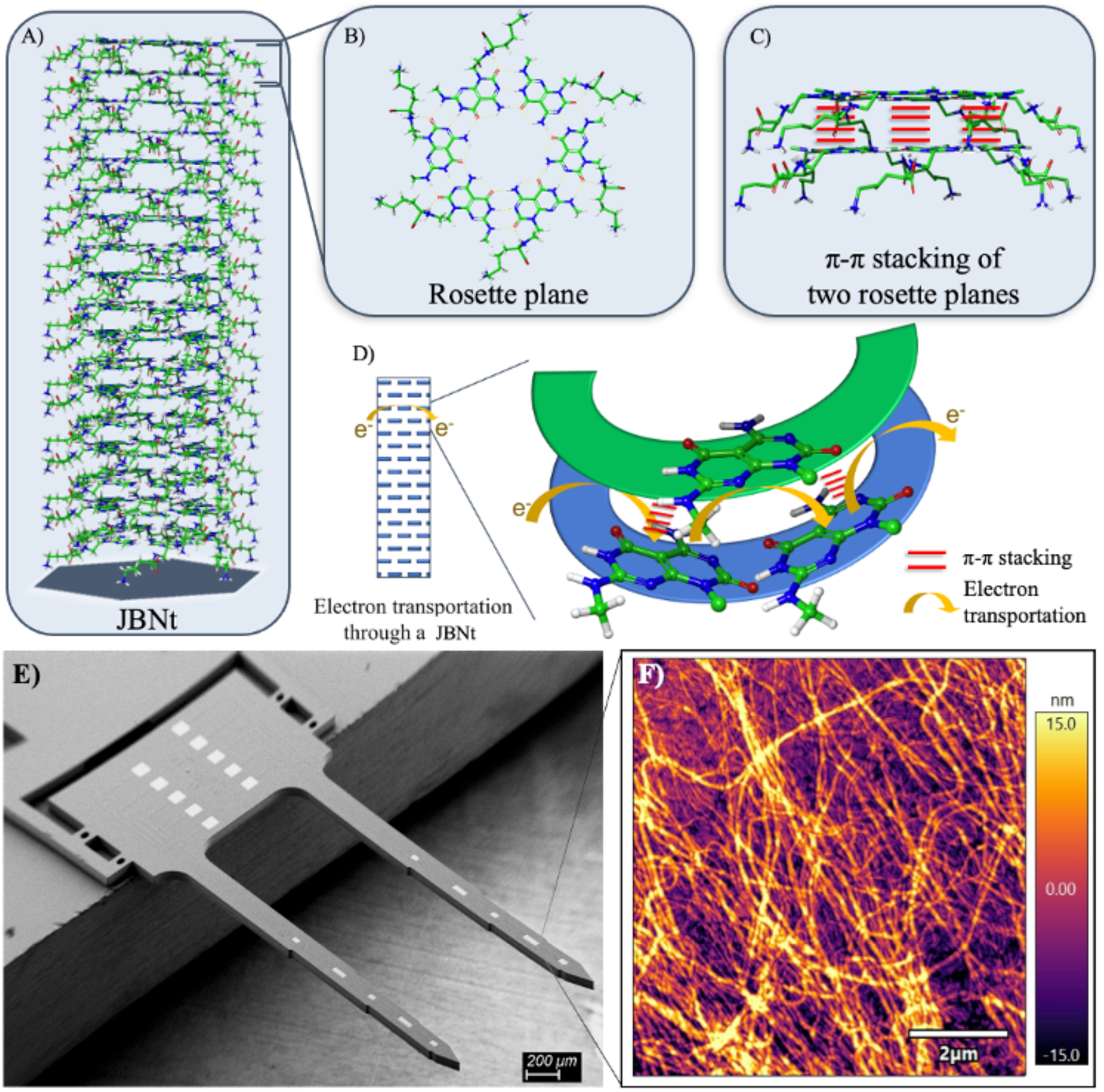
(A) Janus Base Nanotubes (JBNt) self-assembled into long (>100nm) tubes via the base-stacking effect. (B) A superior view of G-C pair orientation stabilized through hydrogen bonding. (C) Stacked rosette conformation expanded to demonstrate electron cloud formation and charge distribution between rosettes in assembled conformation. (D) Electron transport through axis of JBNt assisted by pi-pi stacking. JBNt coating over neural probe devices. An SEM image of a silicon probe with microelectrode sites along the two shanks. AFM images of multiple electrode areas characterizing height and amplitude of JBNc.

### Electrical Conductivity, In Vivo Neural Stimulation and Recording and Elastic Modulus Measurement

The electrochemical properties of the iridium electrodes with and without JBNc were quantified (**Figure 2**). Impedance magnitudes were slightly greater, and phase angles were slightly more capacitive after coating (**Figure 2ai**). Cyclic voltammograms showed that charge storage capacities were similar but exhibited a reduced hydrogen reduction after JBNc were added (**Figure 2aii)**. Most importantly, however, constant-current charge injection pulses produced reduced voltage transients after JBNc addition, which is desired in terms of charge injection capability (**Figure 2aiii)**.

**Figure 2.**
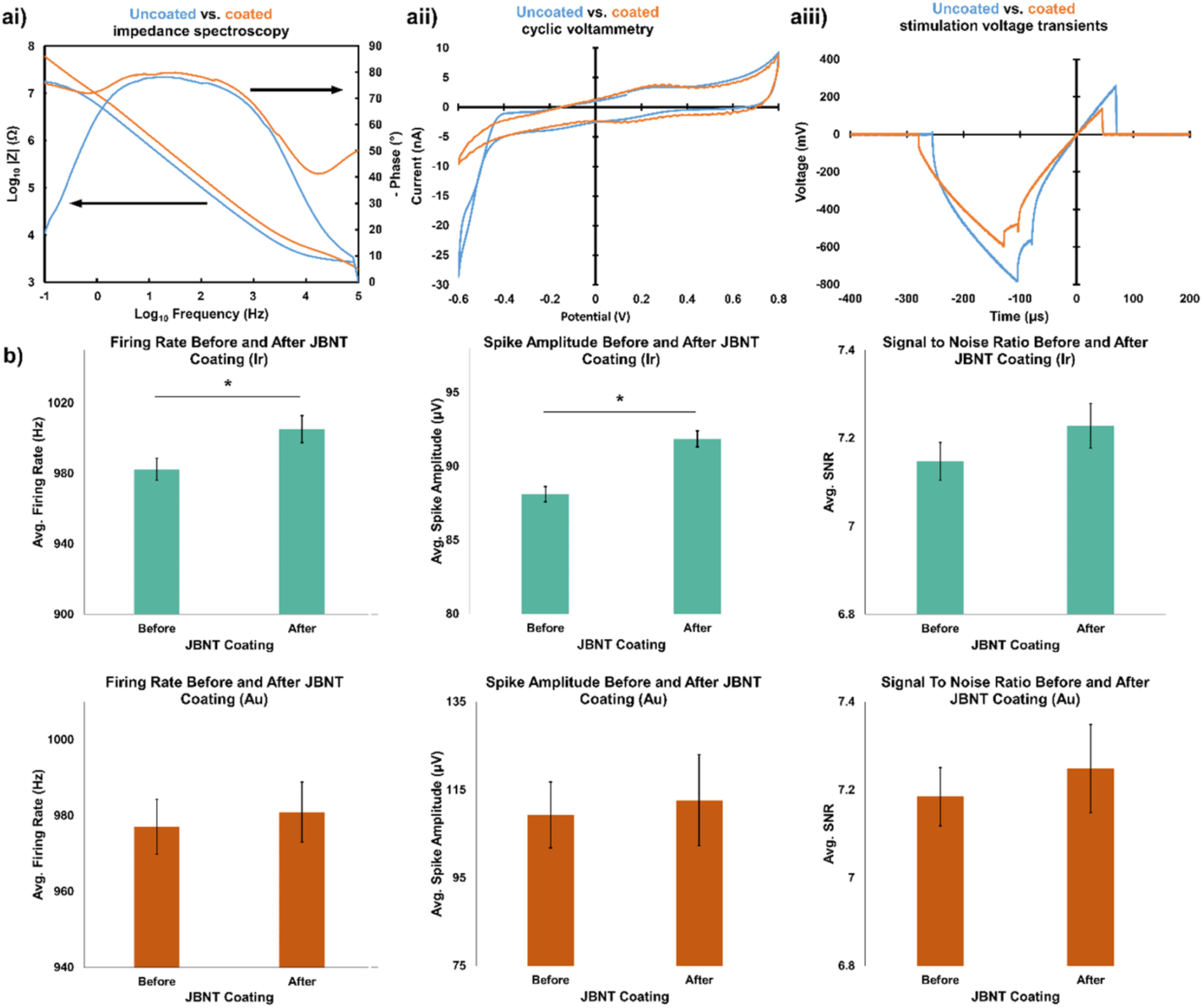
Electrochemical properties of gold and iridium microelectrodes before and after JBNc addition. (ai) Impedance magnitudes were slightly greater and phase angles were more capacitive after coating. (aii) Charge storage capacities were similar, with reduced hydrogen reduction after coating. (aiii) Voltage transient in response to a constant-current pulse was reduced after coating, which is desired. (b) Average firing rate, average spike amplitude, and signal-to-noise ratio for iridium microelectrodes (top row) and gold microelectrodes (bottom row) demonstrated similar or slightly improved capability to detect spike waveforms before and after JBNc addition. (c) Sorted units and high-pass filtered waveforms for iridium (top) and gold (bottom) microelectrodes showed qualitatively similar capability to detect spike activity before and after JBNc addition. *P < 0.05, **P < 0.01, and ***P < 0.001 compared to control.

JBNc had minimal impact on neural probes’ ability to detect simulated neural spike signals *in vitro* before and after JBNc addition (**Figure 2b** and **c)**. Gold microelectrode sites experienced no significant change in measured firing rate, average spike amplitude, or signal-to-noise-ratio (p = 0.73, 0.80, and 0.61, respectively), while iridium microelectrode sites recorded slightly higher firing rates (p = 0.047) and larger spike amplitudes (p < 0.001) after JBNc addition (**Figure 2B)**. No significant change in signal-to-noise ratio was found in iridium microelectrodes, similarly to gold (p = 0.26). Sorted units and high-pass filtered data are illustrated in **Figure 3C** for gold and iridium microelectrodes, which are highly similar to each other before and after JBNc addition.

**Figure 3.**
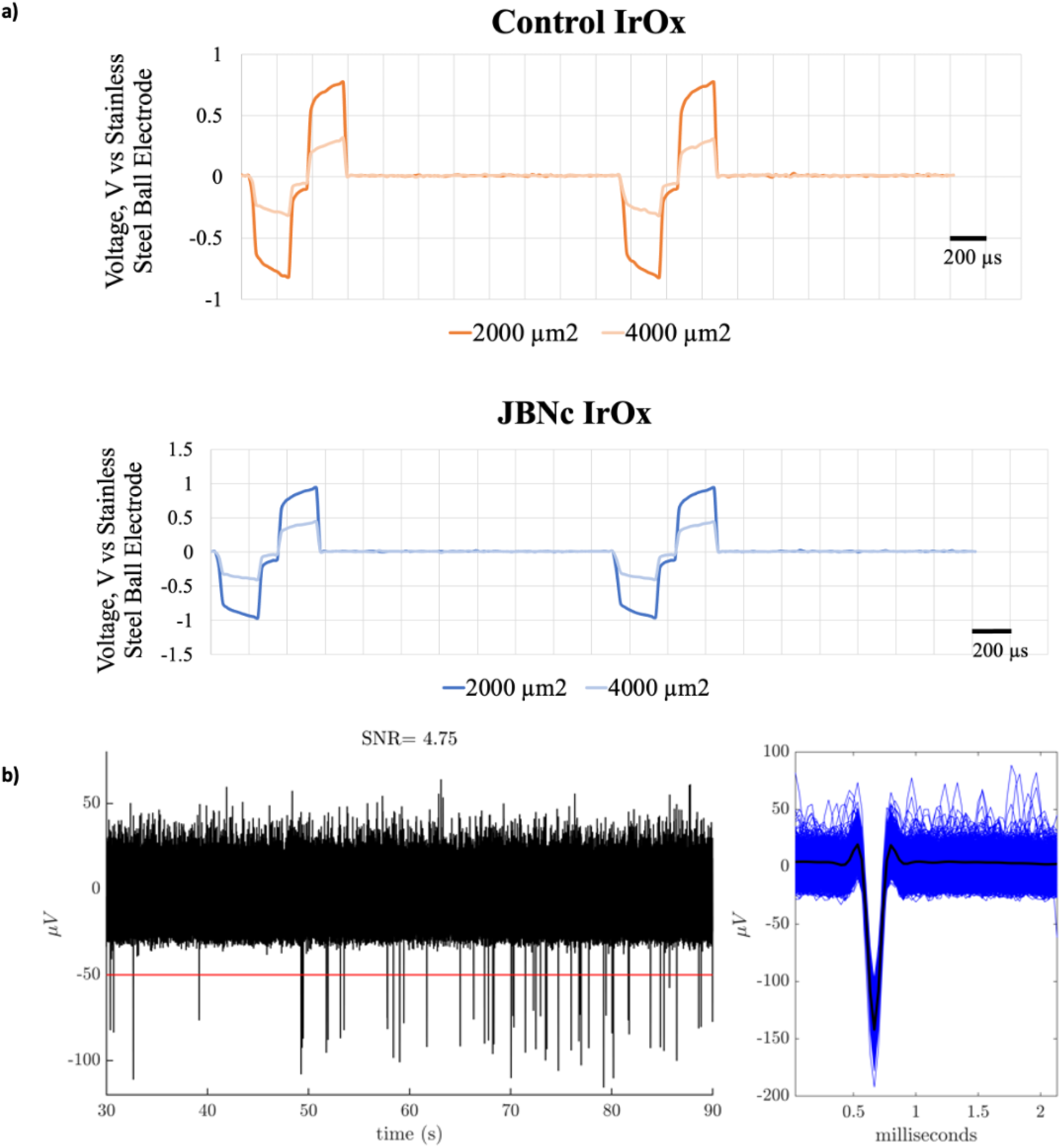
a) Constant-current, charge-balanced, and biphasic waveforms in either 2000 or 4000 µm^2^ electrodes. b) high-pass filtered neural recordings and sorted units taken from JBNc microelectrodes in vivo.

Constant-current stimulation of 10µA, charge-balanced, and biphasic waveforms were injected to the electrodes of either 2000µm or 4000µm areas. In both cases, initial ohmic resistance was higher for JBNc microelectrodes and the level of polarization was comparable compared to IrOx microelectrodes (**Figure 3a**). As it is known that Ohmic resistance does not contribute to the electrochemical damage to the tissue or electrodes, the JBNc was deemed as not inferior to the control IrOx microelectrodes. In the same preparation *in vivo*, high-pass filtered neural recordings and sorted units were taken from JBNc IrOx microelectrode, showing a robust SNR of 4.75 (**Figure 3b**).

Elastic modulus measurements were collected on JBNc SiO_2_ wafers using AFM. JBNc measured 7.1Gpa +/-3.1 GPa, as shown in **Figure 4**. Elastic modulus values were compared to elastic moduli of relevant materials involved in intracortical microelectrode interfacing applications.

**Figure 4.**
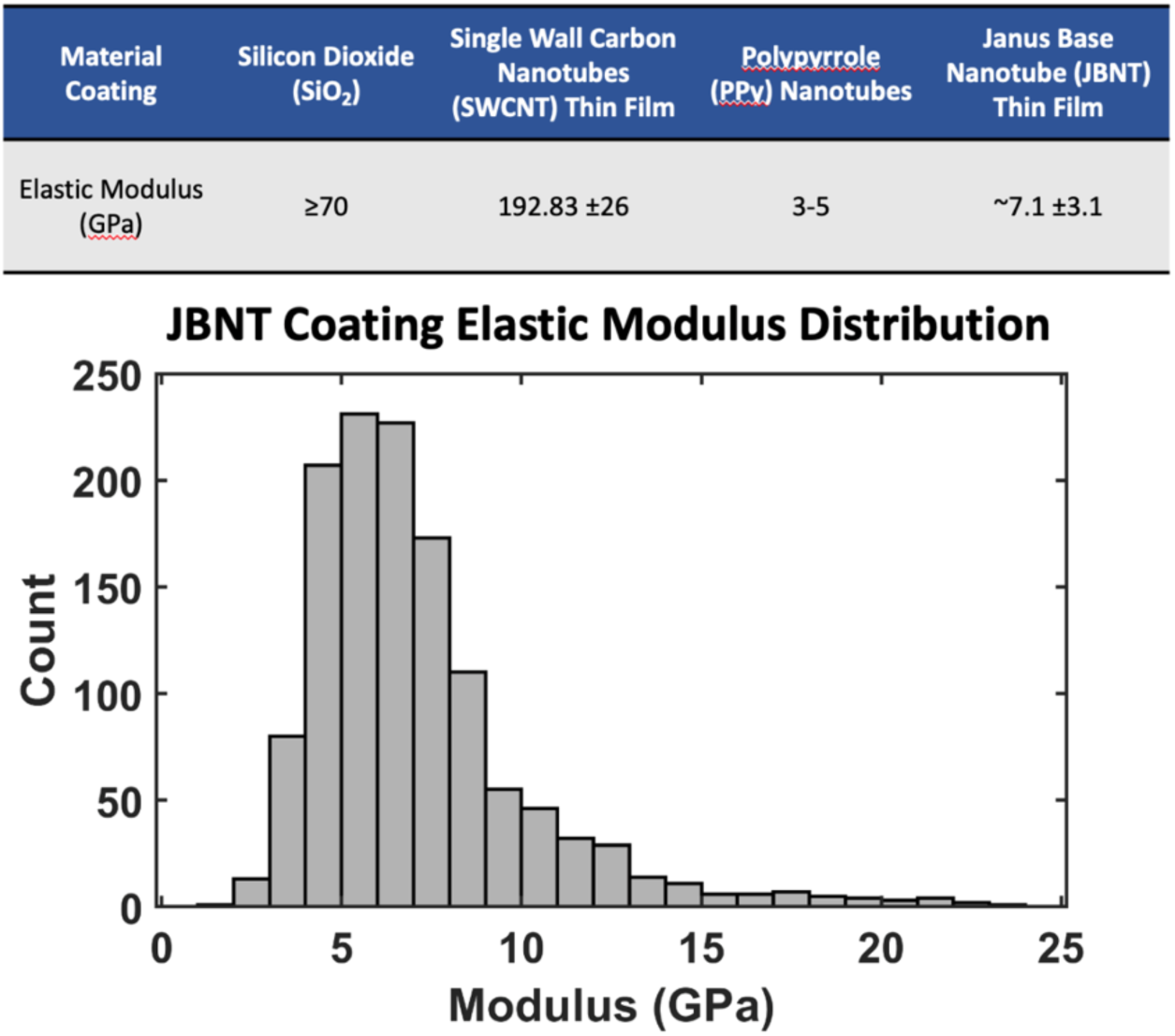
Elastic modulus values summarized from previously published literature. Histogram of elastic modulus measurements of JBNc prepared on coated SiO_2_ wafers.

### Neuron Anchorage, Integration and Functions

SH-SY5Y cells have previously been used to model neuron cell response to CP chemistry and probe modifications[27]. JBNc variables were studied using RSM to optimize neural cell binding and demonstrate a statistically significant linear relationship between JBNc interfacing and neural proliferation after 48hrs *in vitro*. SH-SY5Y cultured on JBNc synthesized at 0.5mg/mL – 1.0mg/mL for >1 coatings produced cell viability of over 1.5X compared to control cells in standard culture conditions **Figure 5a**. RSM results validated the 3X JBNc coating of 1mg/mL used on all electrochemical testing. SH-SY5Y cultured on probes for a 6-day acute timeframe were subject to standard differentiation protocol and subsequently analyzed via confocal microscope to study general cellular behavior patterns. Fluorescence staining showed distinct favorability to JBNt coated probes versus their non-coated control as seen in **Figure 5b**. Increased concentrations of cells were observed to be “crowded” around JBNt coated electrodes as opposed to their control counterparts which demonstrated singular cell activity. Cellular anchorage on and around each electrode site was further analyzed by single cell count. 12-hour and 6-day cultures (respectively **Figures 5c** and **d**) revealed a small favorability towards JBNt coated probes, albeit with high variability between groups. Although a general increase in anchorage was hypothesized for JBNc groups, we predict the large variability is a direct result of *in vitro* well size contrasted with extremely small probe target site for culture. General cell biocompatibility was quantified using two commonly studied CP coatings: single-wall carbon nanotubes (SWCNTs) and Polypyrrole (PPy). CCK-8 assay was used to determine the viability of cells after 24-hour incubation with scaled concentrations of each CP coating material. At lower concentrations (.2-1 μg/mL), PPy presented significantly reduced cell viability compared to JBNt and SWCNTs. At higher concentrations (5-10 μg/mL), JBNt cultured groups maintained statistically significantly improved cell viability in comparison to both conductive materials as seen in **Figure 5e**. JBNts biocompatibility can be largely attributed to the biomimetic composition and hydrogen bonding that produce little to no cytotoxic effect even at relatively large concentrations.

**Figure 5.**
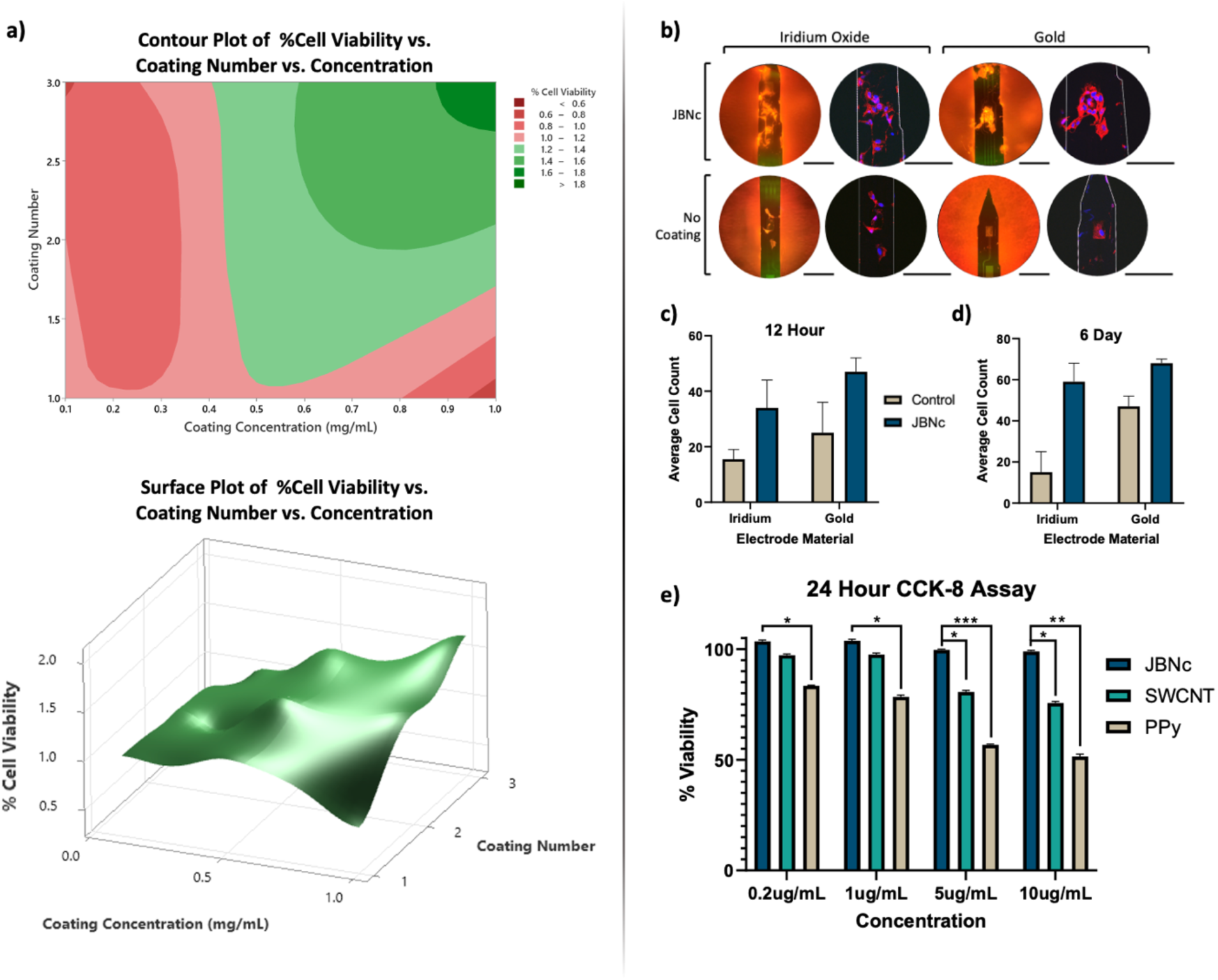
(a) RSM model of SH-SY5Y binding and proliferation on JBNc of varying concentrations and coating numbers (b) Fluorescent microscope (orange) and confocal scan (blue & red) capturing electrode sites after 6-day differentiation of SH-SY5Y cells. Scale bars 100µm. (c) Average DAPI count for iridium and gold electrode probes following 6-day differentiation culture with 50,000 cells/well. (d) Average DAPI count for iridium and gold electrode probes following a short-term 12-hour incubation period with 100,000 cells/well. (e) CCK-8 cytotoxicity assay of JBNt, single-wall carbon nanotubes (SWCNTs), and Polyprryole (PPy) at varying concentrations for 24 hours. Normalized to Control. *P < 0.05, **P < 0.01, and ***P < 0.001 compared to control.

Since JBNc behavior and cell favoritism was consistent between iridium and gold electrode probes, SEM was used to study both probes with respect to cell interface behavior. SEM images were taken to investigate cell-to-electrode interactions as well as neurite advancement onto the JBNt coated substrates fixed after 6-day culture. **Figures 6(a,c,e)** demonstrate typical neural cell attachment on a conventional gold electrode substrate with no additional coating. No cell-to-electrode integration is observed at the nano-scale interface and notable bio-anchorage sites appear at the edge of the electrode well. JBNt coated gold electrodes, on the other hand, showed significant cell-to-electrode interface integration as seen in **Figures 6(b,d,f)**. Higher magnification images further demonstrated cytoplasm outgrowth into nanotube bundle formations. Color overlays for higher magnification images highlight representative boundaries between patches of anchored cells and the electrode surface. These are easily distinguishable for the uncoated control (**Figure 6g**) as opposed to the highly integrated cells (**Figure 6h**).

**Figure 6.**
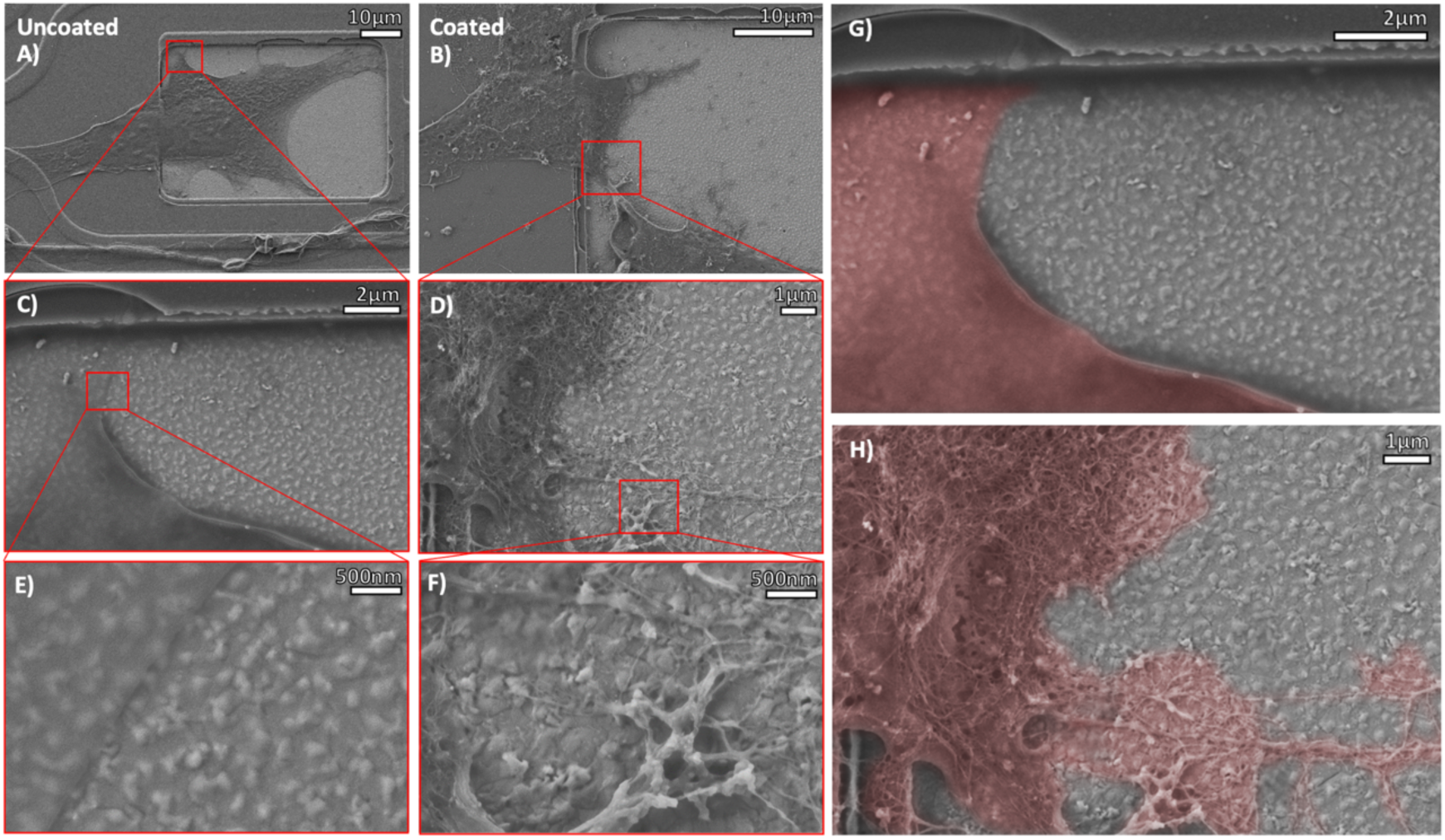
(a,c,e) Qualitative interactions of cell-to-substrate behavior of SH-SY5Y cells to negative control gold electrode with no surface modifications. Scale bars 10μm, 2μm and 500nm respectively. (b,d,f) Cell-to-substrate interactions between SH-SY5Y cells and JBNc Au electrodes sites after 6-day differentiation culture. Scale bars 10μm, 1μm and 500nm respectively. (g,h) Colorized cell dispersion on negative control and JBNt coated electrodes respectively. Scale bars 2μm and 1μm respectively.

Similar SEM imaging also suggests overgrowth between JBNc bundles and neurite progressions for 6-day cultures as seen in **Figure 7a**. Localized AFM scans revealed evidence of bundle formations lying on top of the neurite extensions as seen in **Figure 7b**. We hypothesize that this phenomenon took place during cell proliferation and differentiation as a result of the relatively mechanically compliant biomimetic JBNt bundles.

**Figure 7.**
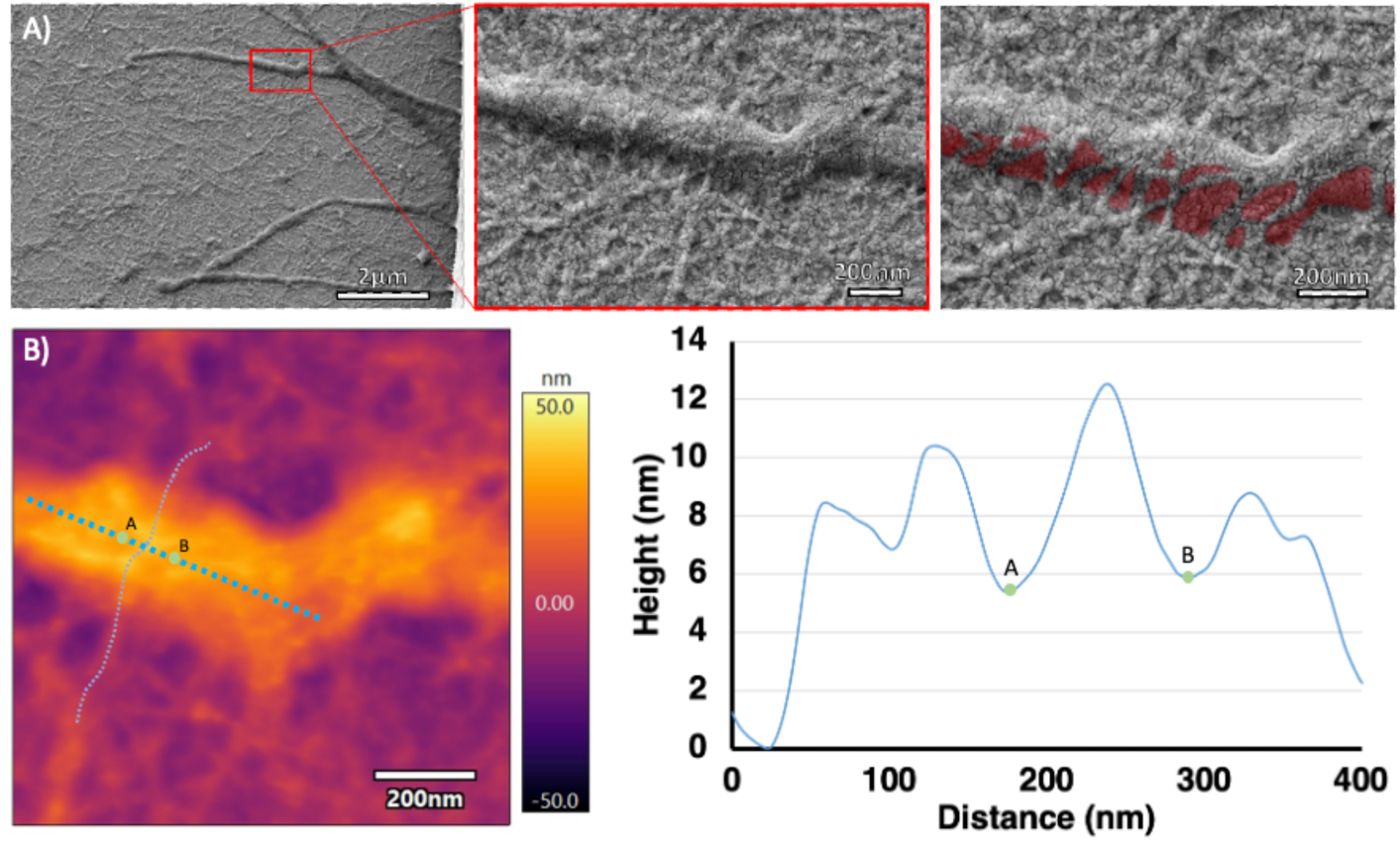
a) JBNc iridium electrode at cell-to-electrode interface. (b) AFM scan of amplitude overlaid onto height of cell-to-electrode interface. Scale bars 2um and 200nm respectively.

It is important to note that presumably due to higher cell anchoring on these optimized probes, interelectrode connections were sometimes observed for JBNc devices. These comprise multiple cell connections terminating on respective electrode surfaces (**S3**). SEM imaging on coated iridium probes cultured long term (14 days) revealed JBNc-integrated anchorage sites remaining after prolonged *in vitro* culture **(Figure 8)**. Complimentary AFM imaging on a separate coated iridium probe (**Figure 8**) suggests that these dense matrices of JBNc withstand prolonged culture conditions to further enhance cell behavior.

**Figure 8.**
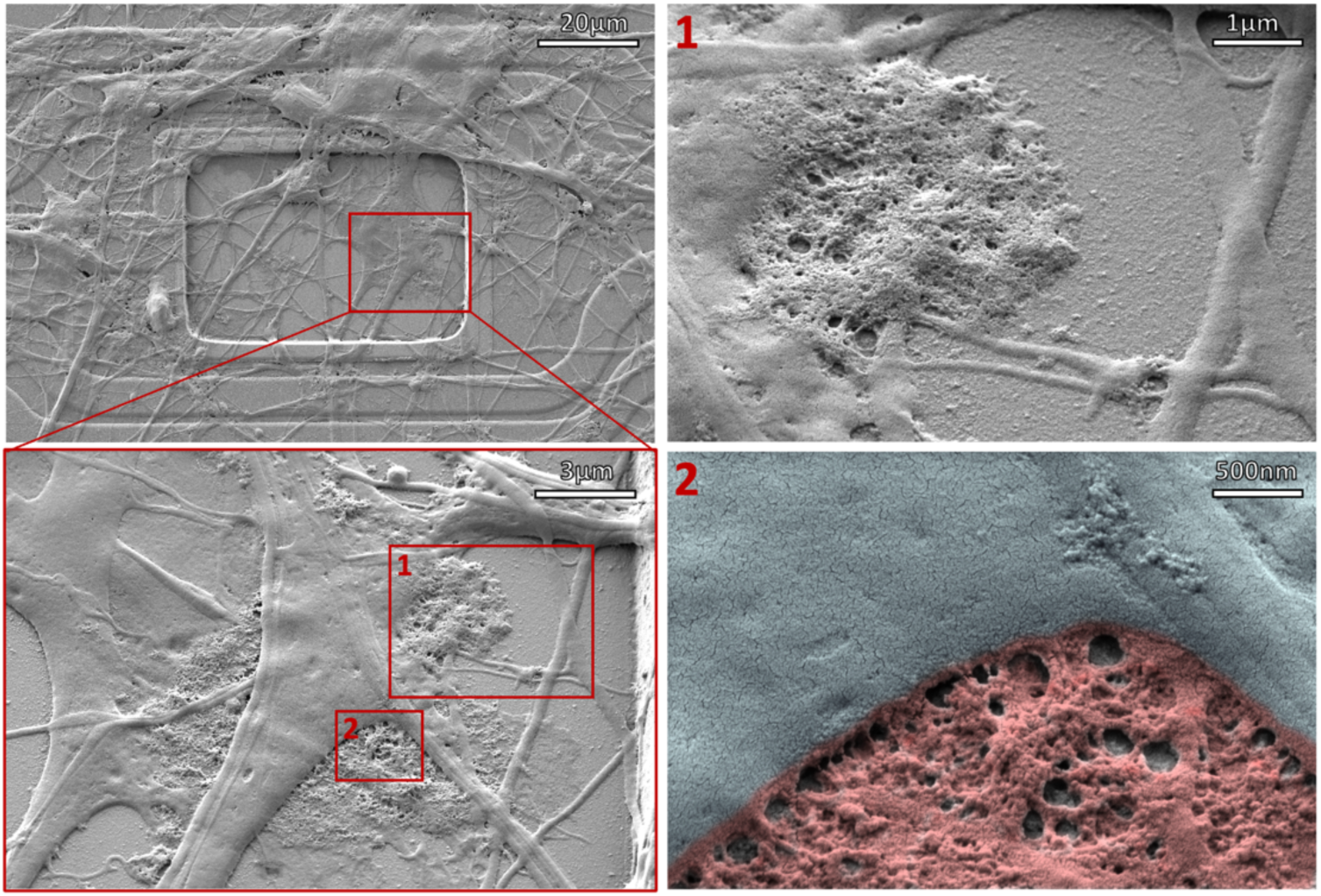
JBNc electrode site following 14-day differentiation culture. High-density areas of JBNt remain and indicate integrated anchorage sites.

Off-electrode qualitative characterization was conducted to determine the selectivity of JBNc as the probes are coated and air-dried. Low magnification JBNc SEM and subsequent regions of interest demonstrated significant JBNc aggregation on multiple material surfaces **(S2)**. JBNc is observed on iridium electrodes as well as the silicon dioxide insulation. Each region of focus investigated the JBNc density and behavior at the electrode, insulation, and cell-to-insulation interfaces. Typical cell integration into the JBNc was observed, further supporting favorable cell attachment behavior. The control demonstrated typical cell behavior observed in previous negative control groups that included rigid cell-to-substrate termination and reduced bio-anchorage **(S2)**.

Immunochemical staining revealed significant discrepancies in cell maturation and growth between experimental groups. β1 integrin staining was conducted to further characterize adhesion performance in the acute timeframe (**Figure 9**). SH-SY5Y cells were deposited onto silicon dioxide substrates with JBNc and collagen 1 fibril coatings and fixed 8 hours later. Single cell and multi-cell regions demonstrate an increase in β1 integrin expression and focal adhesion production during this short adhesion period. Cell surface area and fluorescence intensity increases significantly after coating, as cells on coated surfaces begin to integrate into favorable substrates. Corrected total cell fluorescence (CTCF) analysis on single cell images showed significant fluorescence signal increase in JBNc and Col1 groups.

**Figure 9.**
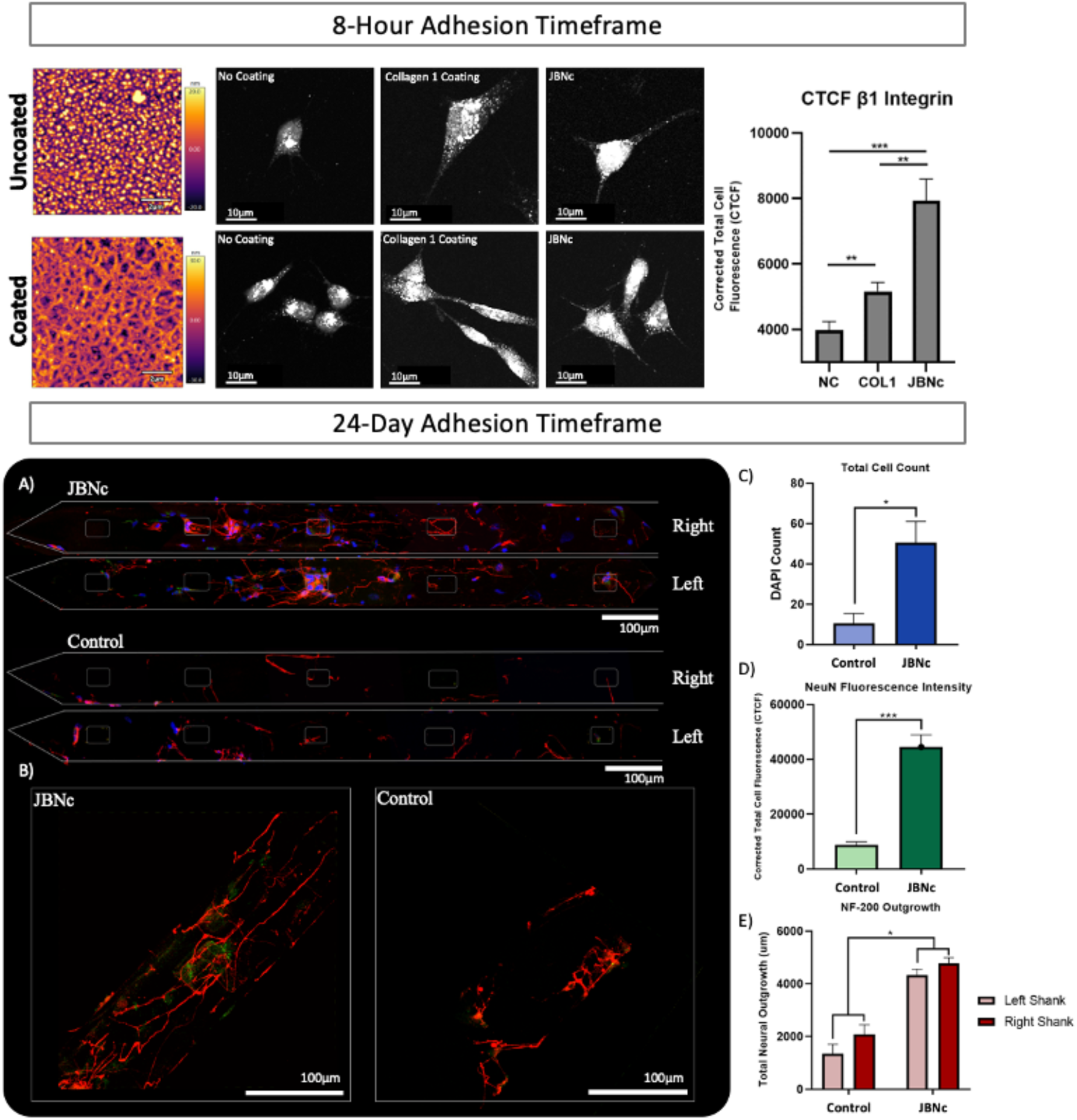
Integrin polyclonal antibody stain on coated silicon dioxide substrates. SH-SY5Y cells were fixed, permeabilized and stained after 8-hour adhesion period to determine acute timeframe focal adhesion behavior. β1 Integrin expression comparable to collagen substrate while both demonstrate superior scaffolding properties in contrast to negative control. Additional CTCF analysis of β1 Integrin expression demonstrating statistical significance. (a) Full shank assembly of JBNc and Control probes triple stained for NF-200 (red), NeuN (green), and DAPI (blue). (b). NF-200 and NeuN strain of most dense electrode sites in each experiment group. (c,d,e) Average cell count, neuronal nuclei corrected total cell fluorescence and neurofilament growth respectively. Scale bars 100µm. *P < 0.05, **P < 0.01, and ***P < 0.001 compared to control.

After two weeks of differentiation culture, sectioned confocal scans were reassembled to show cell coating along the two shanks of each probe (**Figure 9a)**. Representative sections of NeuN and NF-200 expression are shown in **Figure 9b** for comparison. Overall adhesion favoritism was noted by DAPI count which showed a significant difference between groups (**Figure 9c**). NeuN expression on neurons cultured on JBNc probes expressed significantly higher levels of fluorescence compared to the control group (**Figure 9d**). Furthermore, NF-200 staining revealed significantly higher neural outgrowth on JBNc probes (**Figure 9e**). To mitigate any concern of cell seeding density differences, the area surrounding each probe (left, right and center of the shanks) were scanned and cell density was analyzed (**S1**). It was found that although the control shanks demonstrated significantly lower cell count on the regions of interest, the surrounding substrate between and around the control shanks held a significantly higher density of cells in comparison to the surrounding area of the JBNc probe shanks. We believe that this supports the idea that JBNc probes provide a more favorable substrate for cells to adhere to, therefore causing fewer cells to migrate towards the surrounding substrate.

To thoroughly characterize neural cell binding and short-term transcriptomic activity, SH-SY5Y were allowed to culture on JBNc and control SiO_2_ wafter substrates for 72 hours with no neurogenic factor stimulation. These cells were then lysed, and RNA was collected for mRNA sequencing. Following sequence alignment and mapping, cells were grouped based on substrate and analyzed for differentially expressed genes (DEGs) at an FDR <0.1. Differential gene expression was processed in R and analyzed using Gene Ontology (GO)-term and KEGG analysis for statistically significant and differentially regulated gene families present in cells bound to JBNc (**Figure 10**). Many noteworthy patterns arose, including the significant regulation of genes governing focal adhesion/cell-substrate junctions and binding. This paralleled a significant upregulation in cytoskeletal remodeling through microtubule and microfibril activity. Synaptic activity was enhanced during this short culture period, reflected in the upregulation of gene families associated with synapse assembly and neuron projection development.

**Figure 10.**
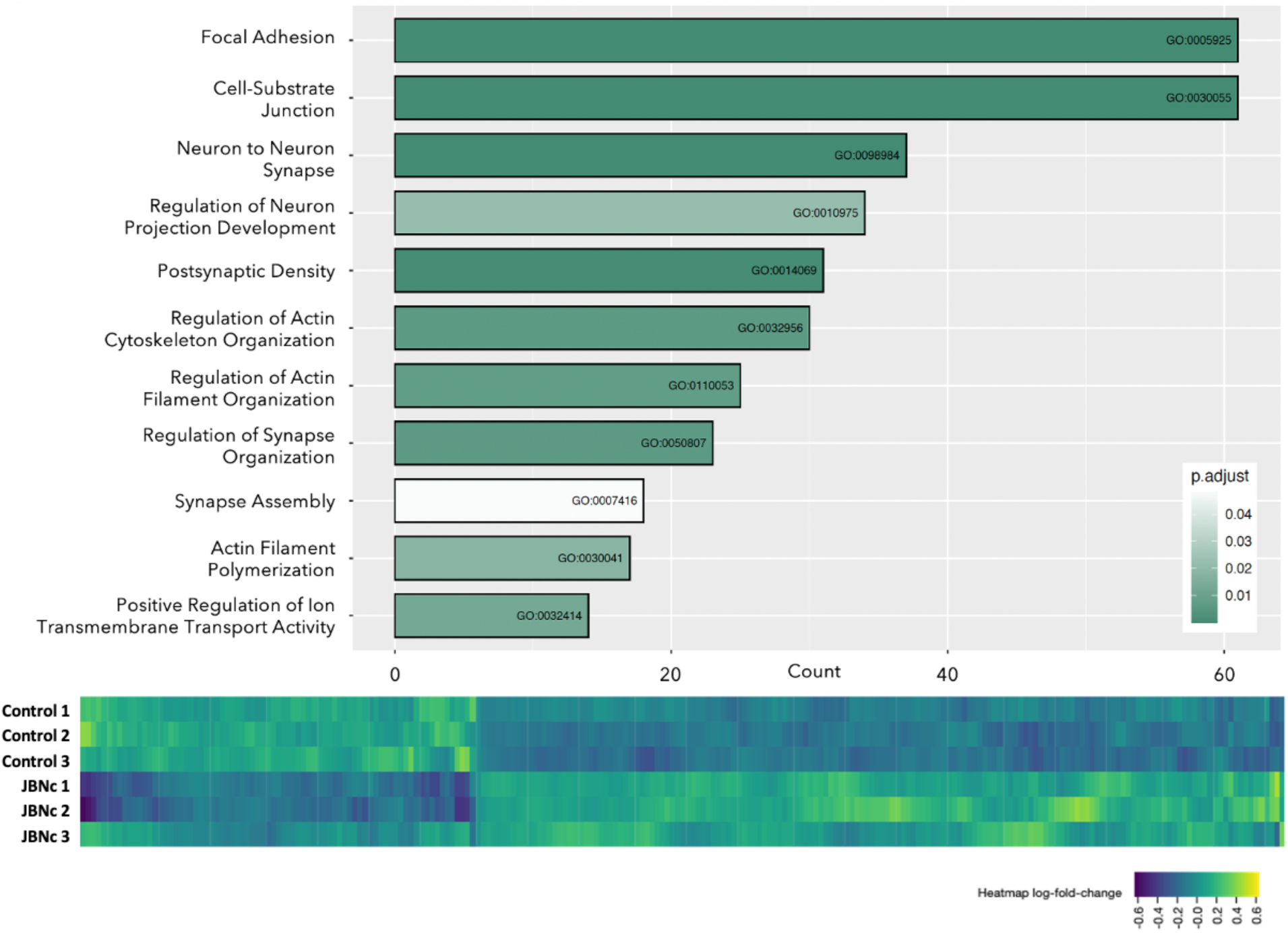
Gene Ontology (GO-term) and DEG heat map analysis of SH-SY5Y cells grown on JBNc versus control. Significant patterns of cell-to-substrate activity are present as well as gene families associated with neuronal differentiation and maturation.

**Figure 11.**
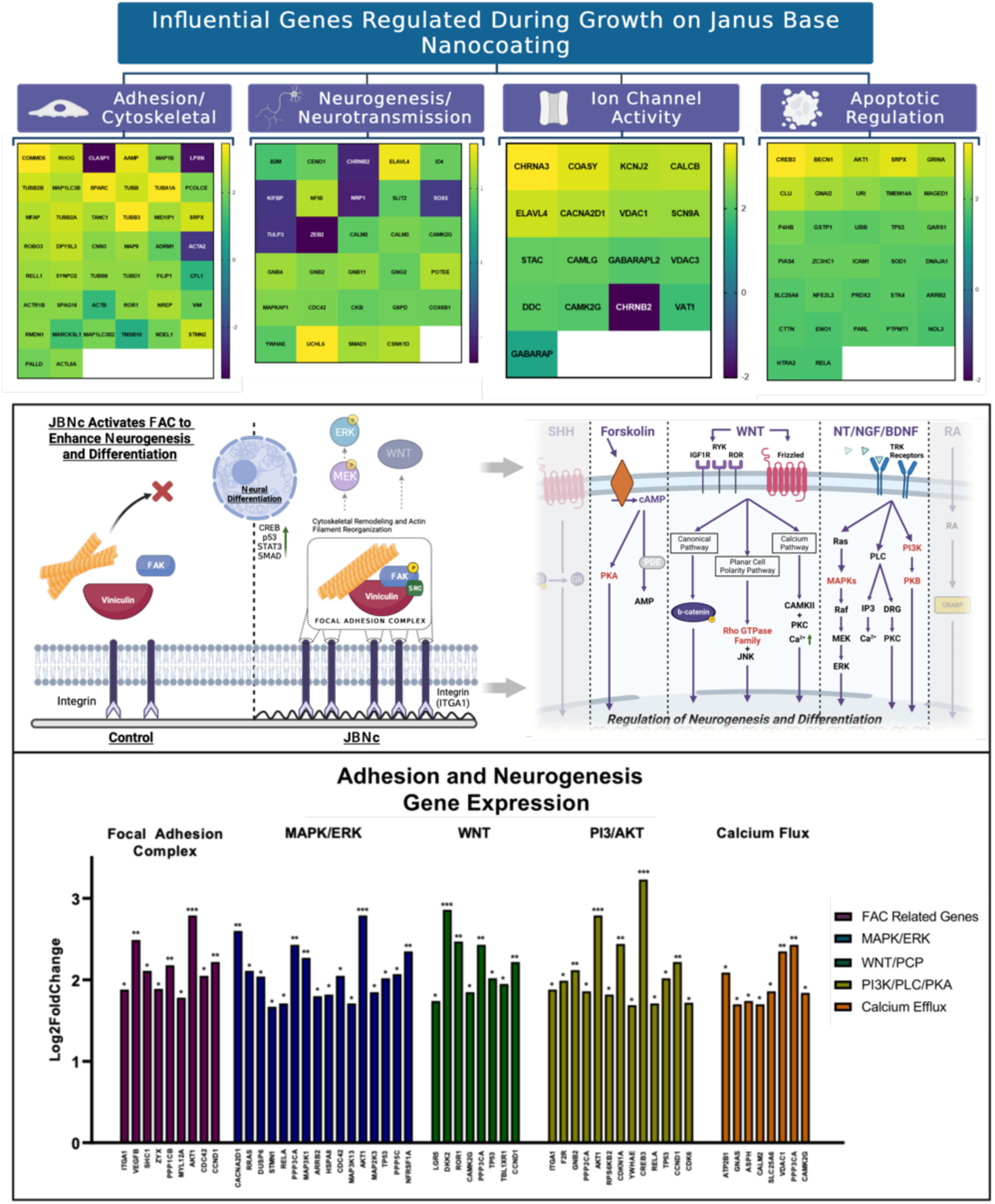
A) Heat maps of influential gene families regulated during neurogenesis and differentiation including adhesion/cytoskeletal rearrangement, neurogenesis/neurotransmission, ion channel activity, and apoptosis regulation. B) Focal adhesion complex (FAC) and associated differentiation pathways augmented by increased FAC activity. C) Genes upregulated during growth on JBNc with respect to differentiation signaling pathways.

Following short-term adhesion analysis, neural cell behavior was studied on probes after an extended differentiation culture on JBNc vs. control shanks to more closely mimic *in vivo* conditions. After two weeks of differentiation culture on JBNc and control microelectrodes, SH-SY5Y cells were manually isolated from their substrate and lysed immediately for scRNAseq analysis. Cells were barcoded and pooled prior to Illumina sequencing and the neural transcriptome for each individual cell was collected and combined by substrate group for analysis. SH-SY5Y differentiation were assessed using groups of genes commonly modulated during neurogenic maturation including cytoskeletal rearrangement, neurogenesis, ion channel activity, and apoptotic regulation. Based on previous evidence of enhanced binding and neurogenic activity, signaling pathways and influential transcriptomic complexes directly effecting neurogenic activity were investigated. The evident enhancement of cell-to-substrate binding and subsequent cytoskeletal rearrangement suggested heavy involvement from the Focal Adhesion Complex (FAC) because of JBNc interfacing. Many genes associated with FAC activity were significantly upregulated in the JBNc group, leading to an increase in FAC-dependent neurogenic processes including proliferation and differentiation.

Due to the varied signaling cascades that may influence neural differentiation, pathways like WNT, MAPK/ERK, PI3K/AKT, and others were studied to identify differentially expressed genes in the JBNc group. As a result of increased FAC activity, cells grown on JBNc were found to express a variety of genes belonging to the MAPK/ERK pathway as well as many belonging to the PI3K/AKT cascade. Additionally, cAMP protein kinase and Rho family GTPase activity were significantly upregulated in cells cultured on JBNc. Although many differentiation pathways demonstrated differential expression as a result of JBNc binding, MAPK/ERK signaling was the most thoroughly activated cascade involved in the enhanced differentiation of SH-SY5Y grown on JBNc.

### Discussion

To evaluate implantable materials, many studies focus on their biocompatibility (such as adhesion and growth of cells on the material surface) while overlook their cytotoxicity (such as viability of cells after they uptake materials debris). For example, CNTs were claimed as biocompatible materials in many studies because cells can adhere and growth on the surface of CNT-based materials without significant abnormality, but it is known that small CNTs are not biodegradable and can cause cell death[28]. This is a major obstacle that limits CNT-based therapeutics for FDA approval. In another example, cobalt metal alloys were considered as biocompatible materials for implantation into patients, but debris generated from these metal alloys was toxic. Ignorance of its cytotoxicity eventually resulted in over $2.5 billion lawsuits[29]. Although a transition to CP-based interfacing electrodes is underway, metal alloys present many significant advantages over current CP technology. CP microelectrode fabrication detracts from the beneficial characteristics of traditional SiO2 insulated microelectrode arrays. SiO2 probes are capable of reducing excessive tissue deformation and dimpling upon insertion which is thought to reduce immune reactions. SiO2 thermal insulation is also a gold standard material to house a high number of small electrodes sites, increasing spatial resolution of surrounding tissue. As a type of implantable materials, electrode coatings can also cause cytotoxicity to neural cells. The impact of cytotoxicity between neural tissue and microelectrode probe increases the production of toxic soluble factors, perpetuating the self-degradation process of the local tissue environment[30]. Today, many types of electrically CP and nanomaterial coatings have been developed, however limited focus has been paid towards the potential for cytotoxic reactions as a result of material debris and cellular uptake. Although many conductive materials can be considered biocompatible due to their ability to interface favorably with neural tissue, reactive oxygen species (ROS) production and cell viability after cell uptake is not often considered. Many of these materials can interface with cells, but inevitably enter the tissue microenvironment and eventually cell cytoplasm. This unintended cell uptake creates significant oxidative stress and ultimately high cell cytotoxicity despite its initial favorable cell interfacing dynamics[31]. A major challenge remains to lessen the reaction to probe implantation by increasing material interface biocompatibility and cytocompatibility in the case of chronic cell uptake. This can be achieved by continuing to develop an electrically conductive biomimetic coating matrix that does not interfere with microelectrode efficacy. Although CNTs and CPs have been thoroughly studied with respects to their toxicological impacts on biological systems, the results have been mixed, with some studies citing relative concentrations and purity to be primary factors in immune response[32, 33]. Although lower concentrations of conductive materials and JBNts present little to no difference in cell viability, relatively higher concentrations of each material demonstrate which candidate can be functionalized as a high-density coating. JBNts remain above 95% viable at a concentration above 10μL while both electrically conductive materials candidates fall below 80%. It is important to recognize that the study of nanomaterial toxicity with respects to CNS interfacing applications is still relatively new, and long-term toxic effects of nano-scale materials are yet to be studied. JBNs possess extremely favorable biocompatibility with various biological systems due to their natural DNA-mimicking chemistry and hydrogen bond-based assembly within stacked rosettes. The absence of carbon bonding during the self-assembly of DNA base mimicking units and natural π-π interactions between rosettes reduces the cytotoxic impact of extracellular and intracellular interactions. Unlike CPs or CNTs, the base units that comprise JBNs are not foreign to the human biological system, leading to tolerability at higher concentrations. This biological familiarity makes JBNt an excellent nanomaterial candidate for long-term microelectrode coating study. JBNc is comprised solely of biological-based nanotubes, making it the first electrically conductive coating of its kind. For the first time, we have demonstrated the conductive potential of JBNc for chronic intracortical microelectrode stimulation and recording.

To serve as a viable coating for neural stimulation and recording applications, JBNc must not negatively impact the electrochemical performance of the microelectrodes on which they are used. Electrochemical characterization (via cyclic voltammetry, electrochemical impedance spectroscopy, and charge injection capacity tests) before and after JBNc addition indicated that JBNc is not detrimental to microelectrode performance, and may even offer some improvement. Charge injection capacity was improved, indicating that JBNc may allow a larger range of stimulation currents to be safely injected. While a slight increase in impedance was observed, this did not appear to negatively impact the microelectrodes’ ability to detect simulated neural signals *in vitro* (a proxy for recording performance *in vivo*). Overall, the results appear to indicate a net improvement to microelectrode performance after JBNc addition.

To further inform the advantageous neural maturation observed on the coated probes, AFM nano-indentation was performed to quantify the elastic modulus profile where cell and probe meet. Current literature has explored the effect of substrate stiffness on neural stem cell differentiation and neurogenesis. Although soft substrates (<1kPa) have be studied as optimal brain-mimicking ECM materials, substrates that exhibit higher elastic modulus values have been seen to enhance voltage gated Ca^2+^ channel currents to increase neural network connectivity[34]. Elastic modulus profiles within the GPa magnitude have been shown to increase directed neurite growth within an unstimulated environment which contradicts the typically ascribed neural environments[35]. Despite its relatively high elastic modulus, SWCNTs are still frequently utilized as coating modifications to neural electrode interfaces. Although SWCNTs increase the electrochemical performance of these modified probes, implantation longevity will remain problematic until bio-integration and neural network integration is achieved. JBNc intracortical microelectrodes exhibited an average elastic modulus of ∼7GPa which is several times less stiff than standard SiO_2_ probe stiffness (≥70GPa) and SWCNT thin film (>190GPa). Although not on the magnitude of native brain tissue, JBNc may act as a softer mechanical biomimetic buffer between foreign electrode material and neural tissue.

On-and-off electrode qualitative imaging demonstrated increased bio-anchorage and enhanced cellular integration onto JBNc microelectrodes. The coated interface has led to notable differences in cell morphology as control group cells maintain an easily distinguishable boundary between cell and surface material, while JBNcs cause more complex cytoskeletal behavior. With respect to shank surface area, nearly 90% of the probe contacting surface is silicon dioxide while the remaining 10% are gold or iridium. Enhanced integration throughout the entirety of the probe is crucial for homogenous biological reaction so silicon dioxide substrates were utilized to study cytoskeletal antigen expression before and after JBNc addition. Quantification of cytoskeletal remodeling was performed with β1 integrin staining on silicon dioxide substrates coated with JBNc and Collagen 1. Previous studies have shown the crucial role β1 integrin plays in neural cytoskeletal remodeling and axon initiation through the homeostatic maintenance of polarization proteins like LKB1 and SAD-A and B[36]. β1 integrin also plays a major role in stabilized microtubule assembly during anchorage and migration during maturation. This data supports the association between β1 integrin production and successful neural anchorage with enhanced differentiation especially on more favorable substrates like JBNc that provide superior scaffolding potential. Two-week monolayer culture onto microelectrodes revealed that neuron cells preferred adhesion to the glass substrate around the control shanks, leading to significantly increased cell density surrounding the target adhesion areas and lower cell counts on those areas of interest. JBNc shanks revealed moderate cell density in surrounding glass areas, far lower than the control group. Furthermore, cell count was significantly higher on those coated shanks, indicating an increase in preferred adhesion. JBNc has also simultaneously enhanced cell maturation as a biproduct of topographical modification[37]. We demonstrated that despite the presence of RA induced differentiation techniques and exogenous growth factor delivery, JBNc cultured cells outperformed control shank cells in NeuN and NF-200 expression. Coated microelectrodes encouraged neurite growth and increased neuronal nuclei antigen production, indicating that JBNc increases mitotic activity and maturation, providing a more favorable platform for neuron function.

Transcriptomic analysis provided a high-resolution dataset that helped illustrate the mechanisms involved during cell-to-JBNc interfacing. Based on short term adhesion studies, cells grown on JBNc displayed in increase in integrin receptor expression due to high aspect ratio and favorable biomimetic chemistry. mRNAseq on 78-hour cultured cells with no neurogenic factor stimulation allowed us to observe the short-term changes driven by JBNc binding alone. Relatively small but significant changes were observed in over ∼200 DEGs which largely contributed to gene families associated with cell-to-substrate junctions and neural cell network formation. Early activity that enhanced axon projection and synaptogenesis were noted through GO Term analysis methods. As we suspected, focal adhesion expression played a significant role in the transcriptomic activity during the 14-day differentiation-induced *in vitro* culture period as well. Many genes involved in the regulation of focal adhesion kinase activity were upregulated in cells grown on JBNc, including the continued upregulation of integrin receptors. Although the FAC does not act as a neural differentiation pathway, its innate function directly contributes to the efficiency of downstream signaling responsible for neural cell function. By increasing FAC activity, downstream cascades such as MAPK/ERK, PI3K/AKT and WNT were enhanced in comparison to cells grown on a control surface under the same culture conditions. Previous studies have studied the complex relationship between cytoskeletal motor proteins and members of the mitogen-activated protein kinase family, suggesting that cytoskeletal stress may have a direct impact on MAPK activity [38, 39]. Additionally, WNT/PCP activity was implicated as a potential pathway utilized during JBNc binding as many Rho GTPases were identified as differentially expressed genes. While some of these signaling cascades were more active than others, it is evident that multiple pathways involved in neural cell differentiation were activated, which illustrates the ability for FAC activity to augment downstream signaling. JBNc acts as a biomimetic scaffolding for prolific cell recruitment and differentiation through FAC-mediated neurogenesis. Due to the significant increase in integration with SH-SY5Y through cell-to-substrate junctions, JBNc enhanced neural differentiation potentially through parallel pathways involving PI3K/ATK, MAPK/ERK, and WNT/PCP.

JBNts possess functional advantages beyond nano-topographical and elasticity modification to neural interfaces, as the hydrophobic hollow core formed within G-C based rosettes may be used for drug encapsulation and delivery[40, 41]. Ionic strength capacity of JBNTs allows for high drug encapsulation and efficiency and strict regulation of drug loading concentrations. JBNts may also be modified with functionalized side-chains to encourage non-covalent bonding to bioactive molecules layered within JBNt bundle formations. Commonly studied anti-inflammatory steroids like Dexamethasone (DEX) have been studied for their modulatory effect on microglial activation and overall immune response suppression, and JBNts have already been proven as a viable platform for slow-release DEX delivery[42, 43]. Antibiotic delivery into electrode implantation sites also demonstrated a reduction in glial scarring as well as a significantly higher signal-to-noise ratio following acute-phase implantation reaction[43]. With these prospects in mind, the development and study of a JBNt-based slow drug release coating is extremely feasible and may assist in lessening the severity of biological inflammatory response directly post-implantation.

## Methods

### Materials and Cell Culture

JBNts were synthesized using an effective approach published previously[44]. Phosphate-buffered saline (PBS, Gibco), Trypsin-EDTA solution (Gibco, 0.25%), Fetal Bovine Serum (FBS, Gibco), Ham’s F-12K (Kaighn’s) Medium (catalog #2112702), EMEM (catalog #302003), DMEM (catalog #11965118), Penicillin-Streptomycin (Gibco, 10,000 U/mL), L-Glutamine (200mM), and Triton X-100 (Invitrogen, 1.0%) were purchased from Fisher Scientific. 4% formaldehyde, 50% Glutaraldehyde, and Ethanol (70% solution) were obtained from Corning. All-trans Retinoic Acid (RA) (R2625) and Brain-derived neurotrophic factor (BDNF human) (B3795) were purchased from Sigma Aldrich.

Human neuroblastoma cell line SH-SY5Y (ATCC CRL-2266) were cultured in 1:1 F12K and EMEM mixture completed with 10% (v/v) FBS at 37°C and 5% CO_2_, with saturated humidity. When cells grew to a density of 70-80%, they were digested with trypsin and passaged in a ratio of 1:4 and were used between passages 3 and 8.

### Microelectrode JBNc

Lyophilized lysine side-chain JBNts were diluted in 1mL of nuclease-free H_2_0 creating a final concentration of 1mg/mL JBNt solution. Non-functional microelectrodes were fixed to 2cm^2^ glass slides using epoxy and placed in 24 well culture plates. 10μL of aqueous JBNt solution was deposited on the surface of the probe, covering the two shanks housing all electrode sites. Probes were air dried and kept in sterile environment until cell culture experiment.

### Cyclic Voltammetry and Impedance Measurements

Electrical Impedance Spectroscopy (EIS) was performed with an Autolab PGSTAT128N potentiostat/galvanostat (Metrohm AG, Herisau, Switzerland) using room-temperature phosphate buffered saline (PBS), a large platinum counter electrode, an Ag/AgCl reference electrode, and 10 mV peak-to-peak sinusoids from 0.1 to 100,000 Hz. Charge injection capability was determined using 150 µs pulse width current pulses from a PlexStim™ stimulator (Plexon, Inc., Dallas, TX, USA), with resulting voltage transients measured with a PicoScope® 2000 USB oscilloscope (Pico Technology, St. Neots, UK).

### Acute *In Vivo* Study

A non-survival study was conducted in order to assess the device’s stimulation and recording properties in the rat motor cortex *in vivo*. The rat was initially anesthetized with 4% inhaled isoflurane, and was monitored using a pulse oximeter and a rectal thermometer for the vital signs throughout the procedure. The rat was mounted in a stereotaxic frame, and was lightly anesthetized with 1%–2% isoflurane during the measurements. The surgical area was shaved and cleaned. A smaller scalpel was used to cut the muscles along the midline. A round spatula was used to lift up some muscle away from the skull and from the target surgical areas. A skull burr hole was drilled approximately 4-5 mm rostral to caudal and 3-4 mm medial to lateral with the rostral end ∼1-2 mm rostral to the bregma. The dura mater was carefully removed to facilitate probe insertion. The microelectrode probe was positioned perpendicular to the brain’s surface, and was successfully inserted (∼2 mm) slowly using the stereotactic tool. A ball-shaped counter electrode was positioned and kept wet on the muscle. The Omnetics connector was connected to the neurostimulator system with a jumping cable, and the ball electrode was connected to the ground of the neurostimulator system. For stimulation, a customized user interface program established the Bluetooth connection between the interface and the neurostimulator. Spontaneous neural recording was conducted using Intan system.

### Probe Culture and Differentiation

SH-SY5Y differentiation protocol used in all 6-day and 14-day culture experiments [45]. Cells were Trypsinized and centrifuged once maintenance cultures reached 70-80% confluency. Cells were evenly deposited within each well at a density between 50,000-100,000 cells/mL. After initial adhesion, cell medium was removed and replaced with Differentiation Media #1, containing DMEM, L-glutamine (4 mM), 1% P/S, and 10μM all trans retinoic acid. Cells underwent RA induced differentiation for 72 hours. Differentiation Media #1 was then removed and replaced with Differentiation Media #2 which contained Neurobasal-A medium (NB) minus phenol red supplemented with 1% (v/v) L-glutamine (200mM), 1% (v/v) N-2 supplement 100x, 1% P/S, and Human BDNF at a concentration of 50 ng/mL. Cells were given 72 hours to culture under these conditions.

### Cell Imaging

After 72 hours of incubation, media was removed, and the samples were rinsed 2x with PBS. For confocal microscopy, cells were fixed with 4% formaldehyde, treated with Triton-X, and stained with Rhodamine Phalloidin (R415, Invitrogen) for 30 min and DAPI (D1306, Invitrogen) for 5 min prior to imagining. Cells were imaged with Nikon A1R Spectral confocal laser scanning microscopy. Images were further analyzed and edited to highlight probe shank outline. All raw DAPI counts were made following fluorescence staining protocol. Immunofluorescence staining was achieved on samples cultured under differentiation protocol and extended for two weeks total. After two weeks, samples were fixed with 4% formaldehyde, treated with Triton-X, and blocked using 10% goat serum (50197Z, Thermofisher) for 30 min. The samples were then incubated in primary antibody solution containing Anti-Neurofilament 200 primary antibody produced in rabbit IgG (N4142, Sigma Aldrich) and biotin-conjugated Anti-NeuN primary antibody produced in mouse IgG (MAB377B, Millipore Sigma) at a uniform dilution of 1:800. A secondary solution of secondary fluorescence-conjugated antibodies containing goat anti-rabbit IgG AF568 (A-11011) and Streptavidin AF488 (S11223). Samples were then preserved using Fluoromount-G (00-4958-02, Fisher Scientific) and allowed to set overnight prior to imaging. Image analysis was performed using corrected total cell fluorescence method and single neuron tracking via ImageJ. β1 integrin staining was performed using the same fixation, permeabilization and blocking method. Samples were then incubated in ITGB Polyclonal Antibody (PA5-29606) produced in rabbit at a 1:500 dilution for 60 minutes. Samples were then incubated in goat anti-rabbit AF488 secondary antibody (A32731) for 60 minutes. DAPI staining was then performed to confirm single cell and multi-cell areas. For scanning electron microscopy (SEM), cells were fixed with 3% glutaraldehyde and treated with a graded series of ethanol dehydrations (50%, 75%, 90%, and 100%). 100% ethanol was allowed to evaporate off samples and each sample was treated with 2nm gold sputter coating prior to imaging. Cells were imaged using ZEISS Crossbeam FIB-SEM for nanometer scale cell-to-substrate interaction characterization. Atomic force microscopy (AFM) images were acquired with a Cypher ES AFM (Asylum Research) and OMCL-AC160TS cantilevers (Olympus) operated in AC mode. JBNt coated electrode surface topography images had 10 μm dimensions with 1024x1024 pixel resolution. Minor variations in height were determined through plotting height profiles averaged across a 50nm wide section. To best observe fixed cells on JBNt coated electrodes, 1 μm images with 1024x1024 pixel resolution were acquired.

### JBNc Elastic Modulus

Elastic modulus measurements of JBNt coated SiO_2_ wafers were collected using a AFM. A. biosphere B100-NCH catilever (Nanotools Bioscience, La Jolla) with a nominal spring constant of 40 N/m and spherical tip radius of 100nm was calibrated and used for all measurements. Force curves were acquired in three 1µm areas using an automated force map to measure 2500 locations per area in a 50x50 array. Each force curve was performed with a trigger force of 2µN and indentation velocity of 2µm/s.

To ensure reported measurements are appropriately measuring bulk JBNt, we utilize the simultaneously acquired height values to establish a mask such that only locations in the top 25^th^ percentile of height values are considered for modulus calculations (i.e. only at the apices of JBNt features). In doing so, we remove concerns of characterizing the underlying substrate instead of the JBNt structures, reduce the likelihood of contact geometry artifacts associated with the edges of the nominally cylindrical features, and ensure our indentations do not exceed 10% of the thickness of the sample as a best practice for any indentation based measurements. The elastic moduli were calculated conventionally using Hertz model fitting with an assumed sample Poisson’s ratio of 0.33[46].

### Cell Toxicity Assay

Cell viability of various materials were determined using CCK-8 assay (Sigma Cell Counting Kit-8, BCCB8166, Sigma Aldrich). SH-SY5Y cells were seeded in 96-well plates at a density of 5,000 cells per well. After 24 hours of attachment, cell media was removed and replaced with media containing various concentrations of JBNt, Polypyrrole, and SWCNTs (.2μg/mL, 1μg/mL, 5μg/mL, and 10μg/mL). After 24 hours of incubation media was removed and cells were gently rinsed 1x with PBS and resuspended in media. 10μL of CCK-8 solution (5mg mL-1) was added to each well and incubated for 2 hours. Absorbance of each well was obtained by SpectraMax M3, Molecular Devices microplate reader at a wavelength of 450 nm. Triplicates were measured for each concentration group and absorbance values, including negative control, were normalized to absorbance of a negative control media and CCK-8 well.

### JBNc Coating Study via RSM

RSM was used to study the relationship of three variables (JBNc concentration, JBNc coating number, and cell viability). RSM was performed in Minitab to produce 3D plots determining the relationship between the independent variables (JBNc concentration and coating number) and the response variable (cell viability).

### Stranded mRNA Sequencing

After the SH-SY5Y cells were cultured on JBNc and control SiO_2_ for 72 hours, the total RNA was isolated using the ___ extraction kits. In brief, mRNA was isolated from total RNA and the mRNA integrity was confirmed using Agilent 1000 TapeStation (Agilent Technologies, Santa Clara, CA). Sequencing was performed on HiSeq 2500 Platform (Illumina) and raw reads were filtered by QC scores <20. The clean reads were aligned to the reference genome using the BWA-MEM function in Galaxy, which maps medium and long reads (> 100bp) against the reference genome. Read count files were merged into a single .csv format and processed through Degust interactive RNA-seq analysis workflow. DEGs were identified using Voom/Limma with CPM set at 0.1 and an FDR cuttoff of < 0.1. To interpret DEG function, Gene Ontology (GO) term analysis and Kyoto Encyclopedia of Genes and Genomes (KEGG) analysis was performed in R.

### Single Cell RNA Sequencing

After the SH-SY5Y cells were cultured on JBNc and control shanks for 2-weeks under differentiation medium conditions, total RNA was isolated using a single cell picking method to transfer individual cells directly into lysis solution. RNA extraction, amplification, purification, and barcoding was performed using the plexWell Rapid Single Cell Kit (seqWell PWSCR96). In brief, mRNA was isolated from total RNA at an average length of 528bp and the mRNA integrity was confirmed using Agilent 1000 TapeStation (Agilent Technologies, Santa Clara, CA). Sequencing was performed on HiSeq 2500 Platform (Illumina) and raw reads were filtered by QC scores <20. The clean reads were aligned to the reference genome using the BWA-MEM function in Galaxy, which maps medium and long reads (> 100bp) against the reference genome. Read count files were merged into a single .csv format and processed through the Degust interactive RNA-seq analysis workflow. DEGs were identified using Voom/Limma with CPM set at 0.1, a >1.5 absolute fold change cutoff, and an adjusted p-value <0.05. To intepret DEG function, Gene Ontology (GO) term analysis and Kyoto Encyclopedia of Genes and Genomes (KEGG) analysis was performed in R.

## Conclusion

We have demonstrated the promising potential that DNA-inspired JBNc holds as a novel biomimetic and electrically conductive coating material for intracortical microelectrodes. Regardless of wet or dry interface properties, JBNc provides robust electrical conductivity to promote and maintain signal transduction and recording. The JBNc matrix increased neuron attachment and enhanced bio-integration and anchorage. NeuN, NF-200 and β1 integrin receptor upregulation was demonstrated on neurons adhered to JBNc substrates, indicating enhanced functional response and maturation. At high concentrations, JBNts presented no cytotoxic effect on SH-SY5Y cells, proving extremely favorable biocompatibility profiles in comparison to CPs and popular nanomaterial coatings. JBNc improved charge injection capacity, and performed favorably in other electrochemical characterizations. Additionally, JBNts are capable of bioactive molecule drug conjugation, which increases their versatility in potential drug delivery applications moving forward.

## Acknowledgements

This study is supported by NIH 7R01AR072027, NIH 1R21AR079153-01A1, NSF Career Award 1905785, NSF 2025362, NSF 2234570, NASA 80JSC022CA006, DOD W81XWH2110274 and the University of Connecticut. BDH and WL acknowledge support from the Institute of Materials Science.

