## Supplemental Data for "Electrically Conductive DNA-Inspired Coating for Intracortical Neural Microelectrodes"

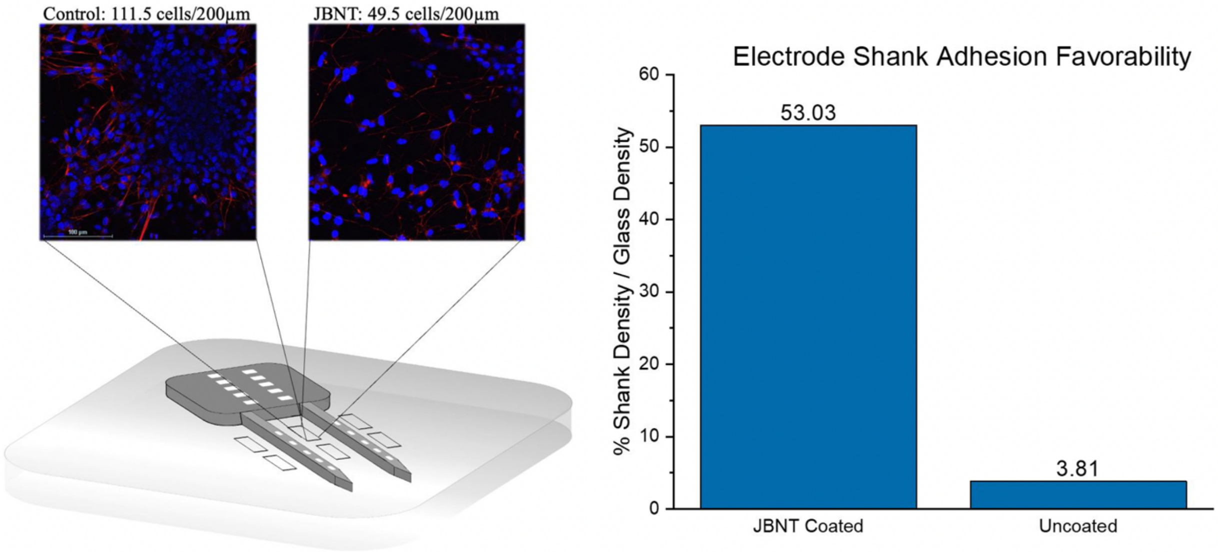


***Supplemental Figure 1*** *Cell density of local culture environment on glass slide is significantly higher in control group versus JBNt group. This suggests substrate favorability is increased in JBNt coated group resulting in less cell repulsion from probe shanks*

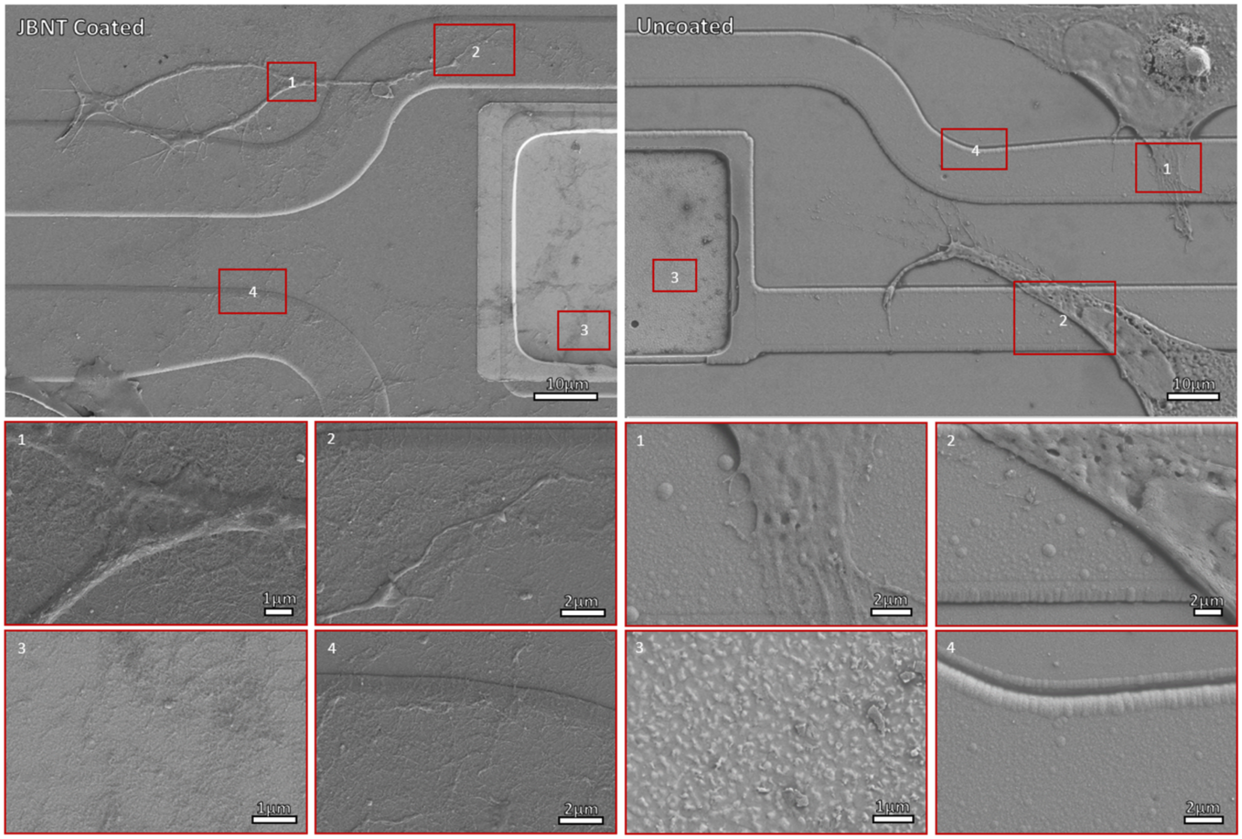


***Supplemental Figure 2*** *SEM scans of iridium and silicon dioxide surfaces with and without JBNt coating. Additional high magnification images (below) demonstrate qualitative differences in surface topography as well as cell interactions with its respective substrate.*

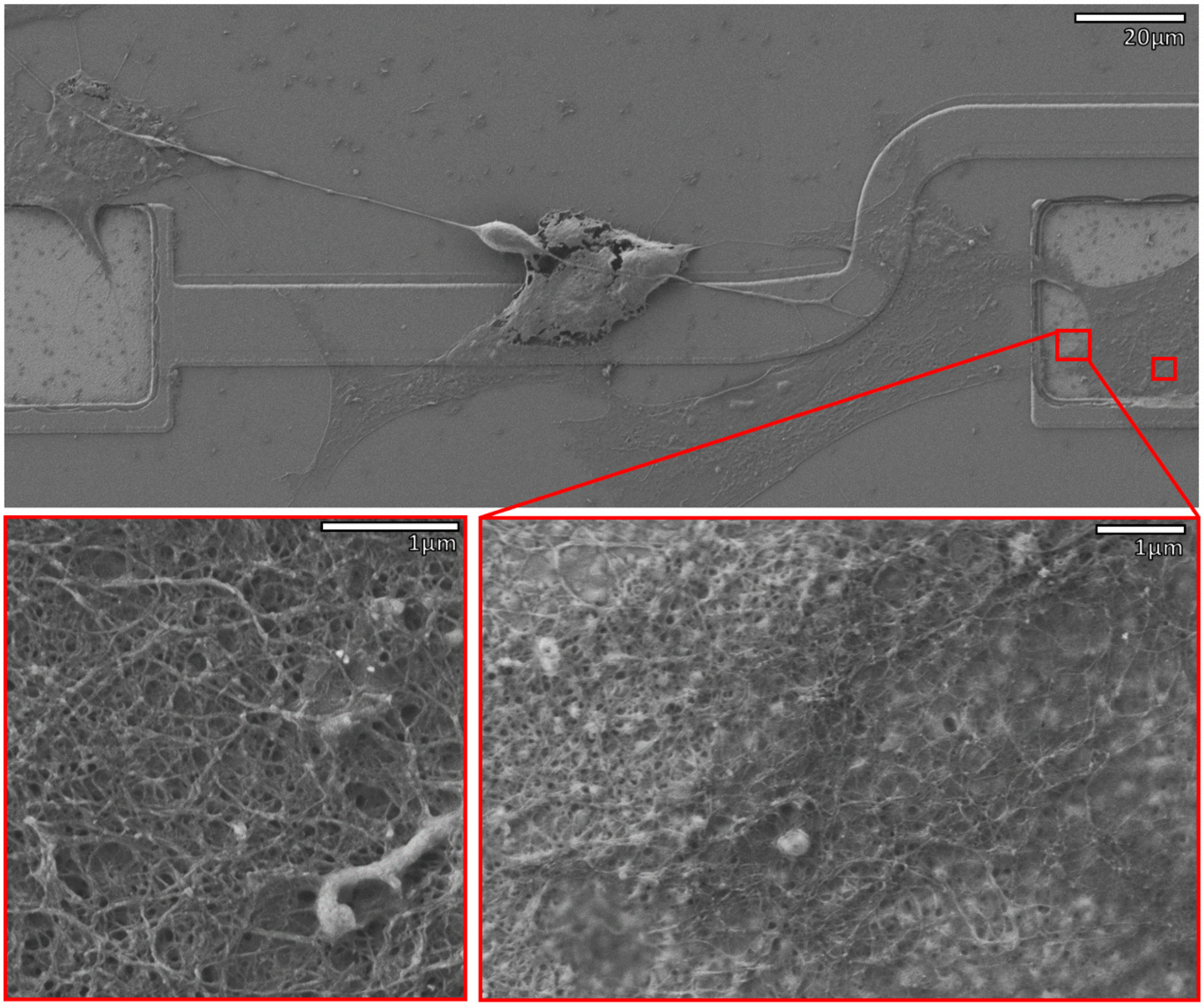


***Supplemental Figure 3*** *SEM scans demonstrating connection between electrodes via cell-to-cell bridging.*

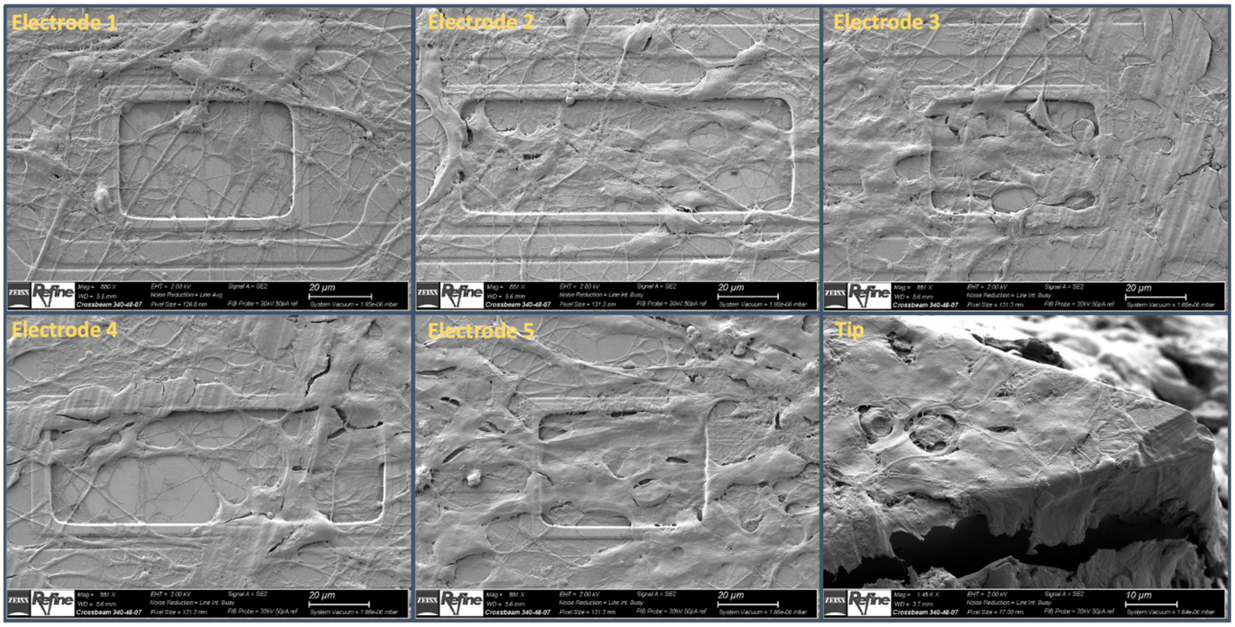


***Supplemental Figure 4*** *SEM scans of select probe shank after 2-week culture on JBNT coated microelectrodes. High cell density noted on both shanks.*

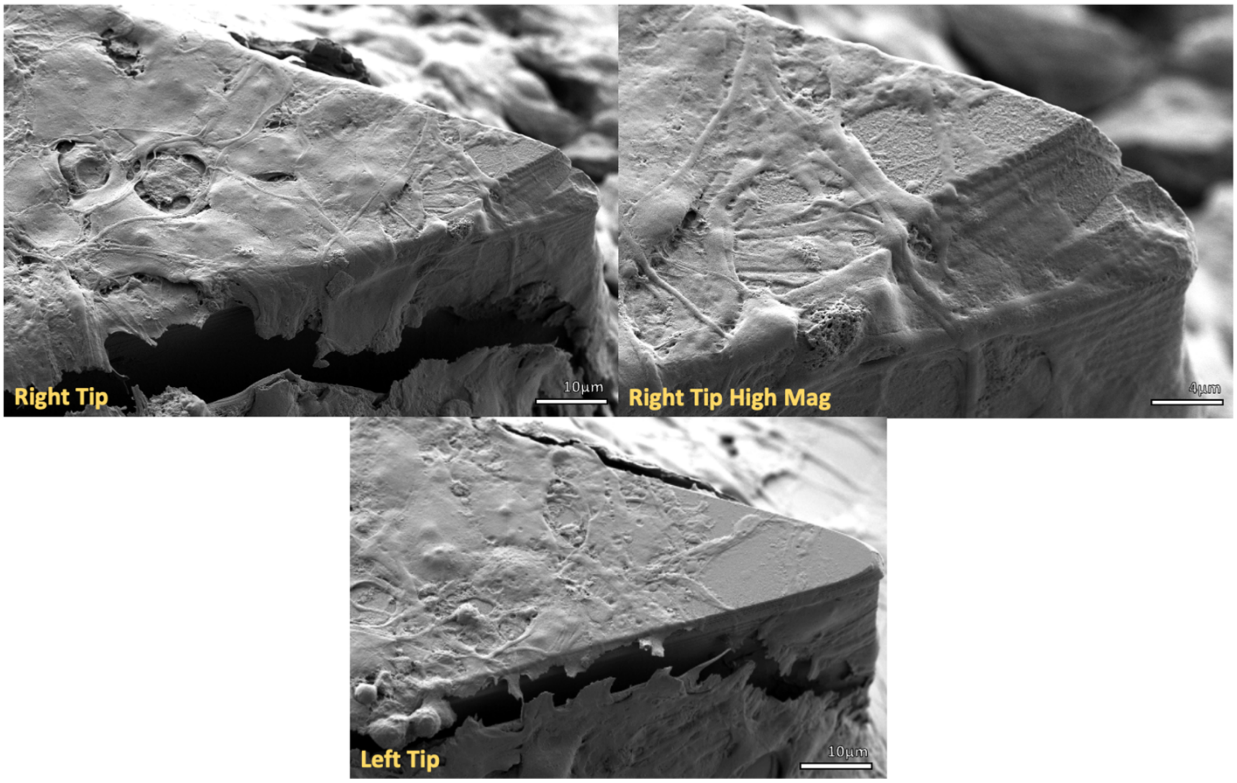


***Supplemental Figure*** *5 SEM scans of probe shank tips coated with JBNt*

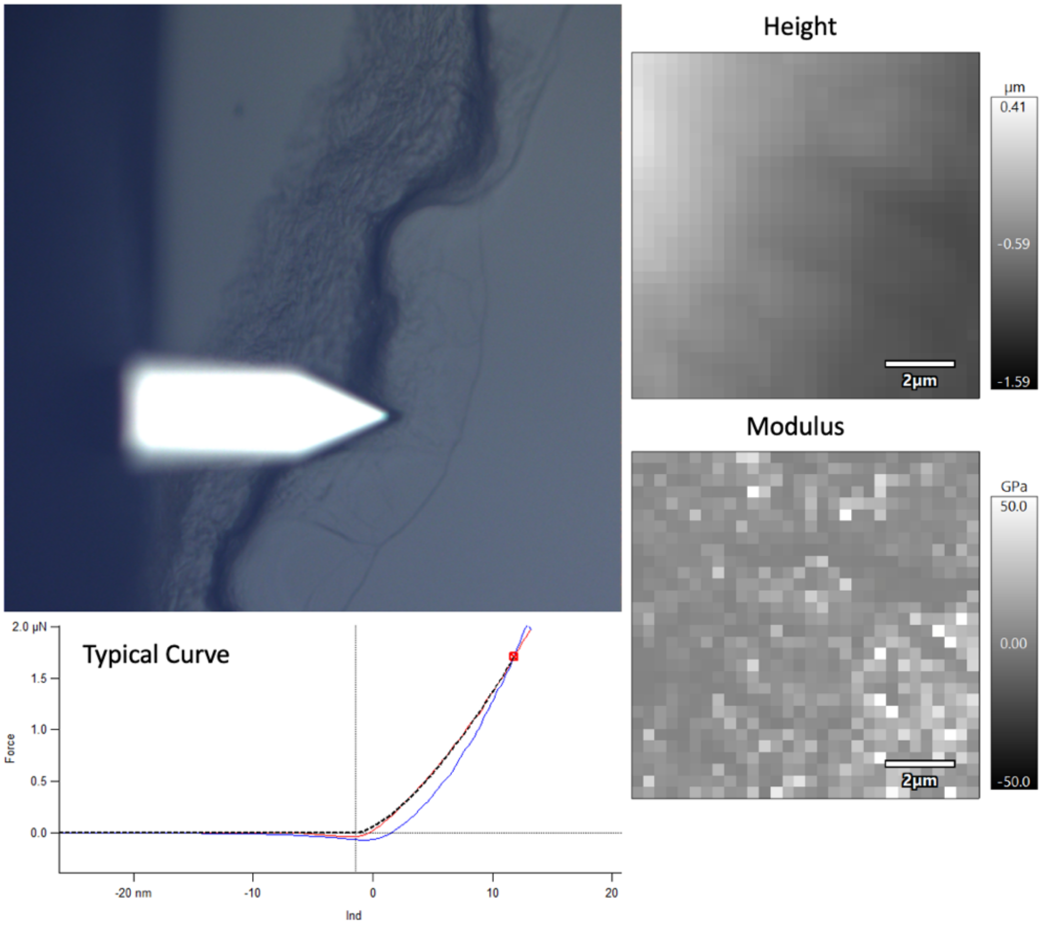


***Supplemental Figure 6*** *Elastic modulus scans and force mapping of JBNt coated $SiO_2$ substrate*

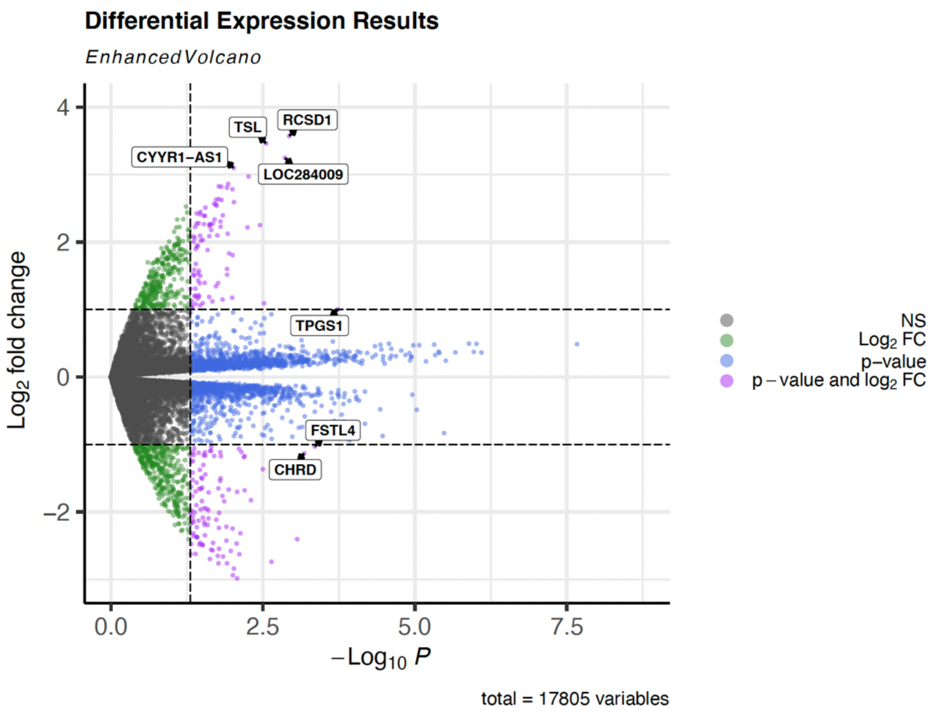


*Supplemental Figure 8 78-hour culture DEGs after JBNc binding (labeled by significance)*

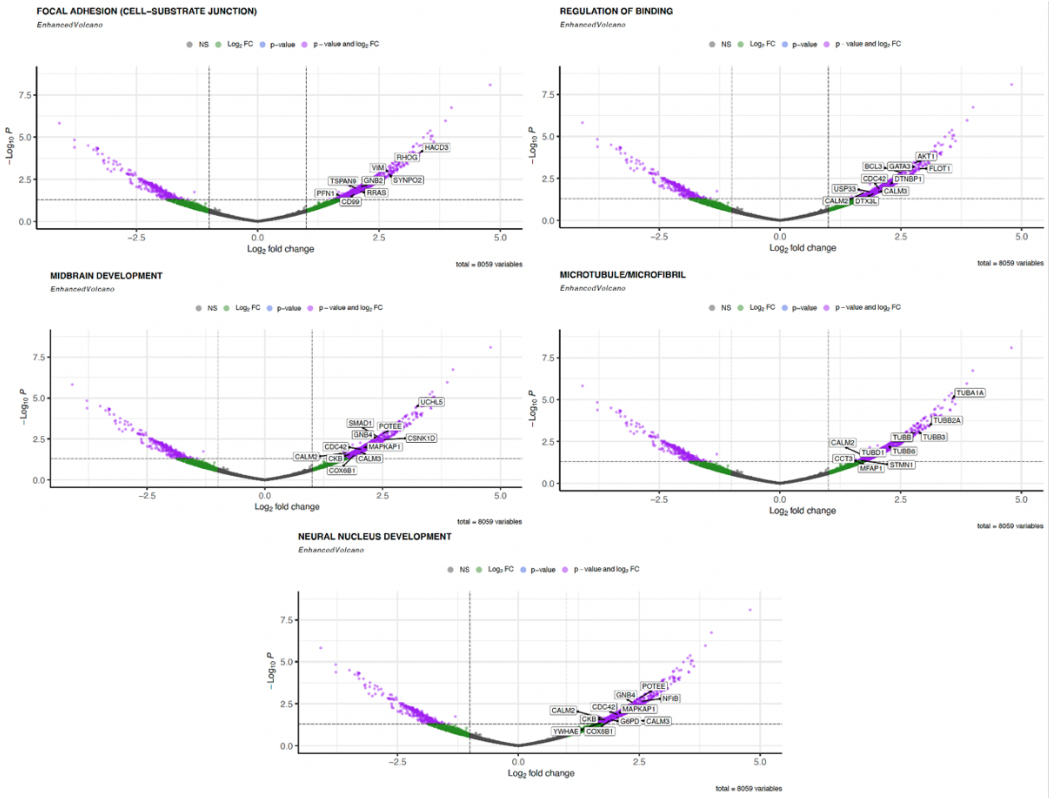


*Supplemental Figure 8 Volcano plots with representative labeling of significantly upregulated GO_Term Analysis Groups*

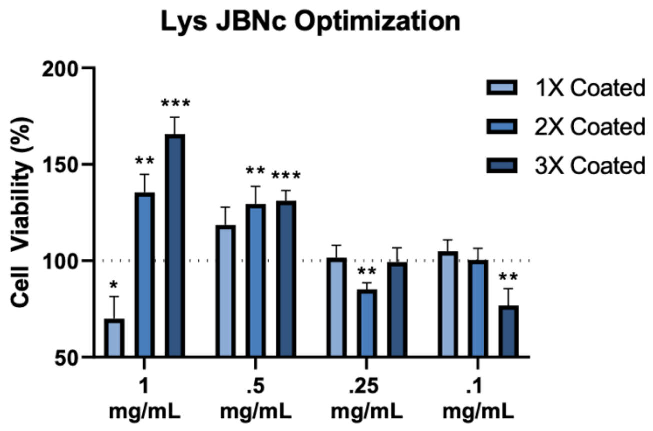


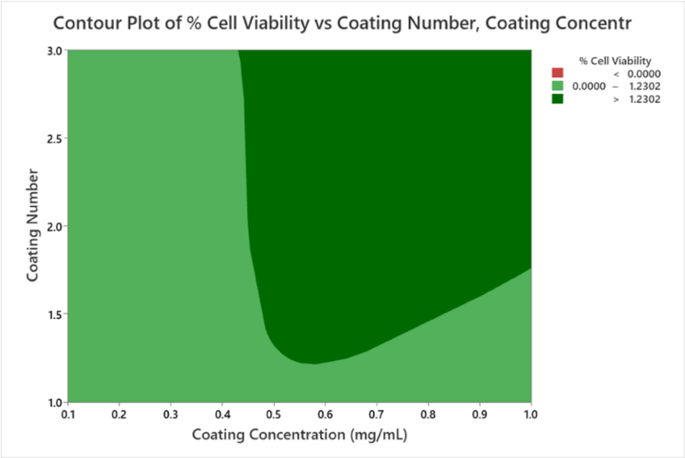


### Coefficients

| Term | Coef | SE Coef | T-Value | P-Value | VIF |
|---|---|---|---|---|---|
| Constant | 0.8207 | 0.0690 | 11.89 | 0.000 | |
| Coating Concentration (mg/mL) | 0.3608 | 0.0919 | 3.92 | 0.000 | 1.00 |
| Coating Number | | | | | |
| 2 | 0.1386 | 0.0769 | 1.80 | 0.075 | 1.33 |
| 3 | 0.1947 | 0.0769 | 2.53 | 0.013 | 1.33 |

### Regression Equation

| Coating Number | | | |
|---|---|---|---|
| 1 | % Cell Viability | = | 0.8207 + 0.3608 Coating Concentration (mg/mL) |
| 2 | % Cell Viability | = | 0.9593 + 0.3608 Coating Concentration (mg/mL) |
| 3 | % Cell Viability | = | 1.0154 + 0.3608 Coating Concentration (mg/mL) |

### Analysis of Variance

| Source | DF | Adj SS | Adj MS | F-Value | P-Value |
|---|---|---|---|---|---|
| Regression | 3 | 2.1013 | 0.70042 | 7.40 | 0.000 |
| Coating Concentration (mg/mL) | 1 | 1.4583 | 1.45826 | 15.40 | 0.000 |
| Coating Number | 2 | 0.6430 | 0.32150 | 3.40 | 0.038 |
| Error | 92 | 8.7108 | 0.09468 | | |
| Lack-of-Fit | 8 | 4.5117 | 0.56396 | 11.28 | 0.000 |
| Pure Error | 84 | 4.1991 | 0.04999 | | |
| Total | 95 | 10.8121 | | | |

***Supplemental Figure 9*** *Bar graph representation of data collected for RSM. Confidence interval contour plot for cell viability vs. coating number vs. coating concentration. Model results and statistics with statistical significance.*
